# Post-stroke dendritic arbor regrowth – a cortical repair process requiring the actin nucleator Cobl

**DOI:** 10.1101/2021.08.26.457764

**Authors:** Yuanyuan Ji, Dennis Koch, Jule González Delgado, Madlen Günther, Otto W. Witte, Michael M. Kessels, Christiane Frahm, Britta Qualmann

## Abstract

Ischemic stroke is a major cause of death and long-term disability. We demonstrate that middle cerebral artery occlusion in mice leads to a strong decline in dendritic arborization of penumbral neurons. These defects were subsequently repaired by an ipsilateral recovery process requiring the actin nucleator Cobl. Ischemic stroke and excitotoxicity, caused by calpain-mediated proteolysis, significantly reduced Cobl levels. In an apparently unique manner among excitotoxicity-affected proteins, this Cobl decline was rapidly restored by increased mRNA expression and Cobl then played a pivotal role in post-stroke dendritic arbor repair in peri-infarct areas. In *Cobl* KO mice, the dendritic repair window determined to span day 2-4 post-stroke in WT strikingly passed without any dendritic regrowth. Instead, Cobl KO penumbral neurons of the primary motor cortex continued to show the dendritic impairments caused by stroke. Our results thereby highlight a powerful post-stroke recovery process and identified causal molecular mechanisms critical during post-stroke repair.

## Introduction

Five million people remain permanently disabled after stroke each year. In the infarct area, stroke leads to a loss of neurons and neuronal network connections due to lack of energy, excitotoxicity, oxidative stress, inflammation and apoptosis as pathophysiological events (Hossmann, 2006). Ischemic stroke caused by middle cerebral artery occlusion (MCAO) accounts for approximately 70% of all infarcts (Bogousslavsky et al., 1988; Musuka et al., 2015) and can also be achieved experimentally in rodents (Fluri et al., 2015). The relative lesion size of survivable human stroke is usually limited to a few percent of the brain (Cramer et al., 2006). In mice, such damages are very well resembled by 30 min induced MCAO. In contrast, prolonged paradigms do not resemble survivable human strokes, as they lead to a loss of large parts of the entire hemisphere affected and to significant structural changes at distant or even contralateral sites (Cramer et al., 2006; Winship and Murphy, 2008; Popp et al., 2009).

Ischemic stroke does not only lead to neuronal death and loss of connectivity inside of infarct areas. It also causes a loss of synapses (dendritic spines) adjacent to the infarct area (penumbra) (Corbett et al., 2006; Brown et al., 2008; Wu et al., 2014). The spine loss is associated with excessive Ca^2+^ influx activating numerous downstream effects. Prominent among those is the Ca^2+^-activated calpain-mediated break-down of synaptic scaffold and receptor proteins, such as spectrin/fodrin, PSD95 and NR2B (Simpkins et al., 2003; Pike et al., 2004; Gascón et al., 2008). Importantly, in the penumbra, these synaptic defects are transient and are subsequently repaired, as spines remain somewhat plastic even during adulthood (Koleske, 2013). Dynamics of dendritic spines (also referred to as dendritic remodeling, dendritic plasticity or synaptic plasticity) is considered as important cellular mechanism for learning but also for compensatory synaptic repair subsequent to stroke (Brown et al., 2007; Brown et al., 2008; Wu et al., 2014; Murphy and Corbett, 2009; Di Pino et al., 2014; Jones, 2017).

Most dendrites and dendritic branches become stabilized at an age of about 20 days in mice (Koleske, 2013) and much less is known about the fate of the dendritic arbor in the penumbra during the acute phase of cerebral ischemia in mice when compared to dendritic spine dynamics. The reason is that the majority of published studies focused on putative long-term effects inside and outside of ischemic areas several weeks or even several months after (often massive) brain damage caused by cerebral ischemia induced by different means in rats (Gonzalez and Kolb, 2003; Brown et al., 2007; Brown et al., 2010; Garcia-Charvez et al., 2008; Winship and Murphy, 2008; Mostany and Portera-Cailliau, 2011; Nesin et al., 2019). Other studies tried to identify long-term effects at the contralateral side that may represent compensational processes (Biernaski and Corbett, 2001; Biernaski and Corbett, 2004; Papadopoulos et al. 2006) rather than focusing on the ipsilateral penumbra neighboring the infarct area.Recent reports suggested that inside of the penumbra, acute changes in the dendritic arbor ocurr or can be pharmacologically induced (Hu et al., 2020; Mauceri et al., 2020), respectively.

The morphology of the complex dendritic arbor of neuronal cells is stabilized by microtubules, yet, it is the *de novo* generation of actin filaments that powers the formation of initial protrusions from dendrites that establishes the dendritic arbor (Kessels et al., 2011). The Wiskott-Aldrich domain 2-based actin nucleator Cobl (Ahuja et al., 2007; Qualmann and Kessels, 2009) (gi:32251014) is widely expressed in the brain (Haag et al., 2012) and was demonstrated to be critical for dendritic branching of developing hippocampal neurons (Ahuja et al., 2007) and of Purkinje cells (Haag et al., 2012) together with assessory machinery (Haag et al., 2012; Schwintzer et al., 2011; Izadi et al., 2021). Interestingly, the functions of the actin nucleator Cobl hereby show regulations by arginine methylation by PRMT2 (Hou et al., 2018) and by multiple Ca^2+^/calmodulin (CaM)-mediated mechanisms directly converging onto Cobl (Hou et al., 2015).

Here we show that induction of ischemic stroke in mice leads to a rapid degradation and a subsequent reexpression of the actin nucleator Cobl. Cobl degradation is excitotoxicity-mediated. By analyzing *Cobl* KO mice (Haag et al., 2018), we furthermore demonstrate that Cobl is crucially involved in a process of regrowth of the dendritic arbor, which we unveil to occur in the penumbra in a narrow time window from day 2 to day 4 after ischemic stroke. With Cobl-dependent post-stroke dendritic arbor regrowth, our work adds a powerful cell biological process inside of peri-infarct areas that represents an acute and long-range mechanism of post-stroke repair.

## Results

### Degradation of the actin nucleator Cobl upon MCAO

The actin nucleator Cobl (Qualmann and Kessels, 2009; Ahuja et al., 2007) is regulated by Ca^2+^/CaM at physiological Ca^2+^ levels (Hou et al., 2015). Excitotoxicity situations, such as massive neurotransmitter releases upon stroke, lead to pathophysiologically high Ca^2+^ levels. Strikingly, quantitative biochemical analyses of brain lysates of mice subjected to ischemic stroke by MCAO (Fig. 1A-P) showed a reduction of Cobl protein levels at the ipsilateral side. In comparison to intrinsic contralateral controls, ipsilateral Cobl protein levels declined by about 25% 3 h and 6 h after MCAO (Fig. 1A,B,E,F). Unchanged levels of β3-tubulin demonstrated that the observed Cobl loss did not merely reflect neuronal degradation in the stroke-affected area of the striatum per se. Instead, the effect seemed to represent a wide-spread targeted degradation of the actin nucleator Cobl (Fig. 1A-D,I-L). In line, the levels of an excitotoxicity-induced 140 kDa spectrin/fodrin fragment (Lynch and Baudry, 1984; Siman et al., 1984) increased upon MCAO. Similar to the Cobl decline, also the MCAO-induced spectrin/fodrin degradation showed a fast onset (Fig. 1A-D,M-P). The ischemia-induced 140 kDa spectrin/fodrin fragment was still found to be significantly elevated even at 24 h after MCAO (Fig. 1O).

**Fig. 1.**
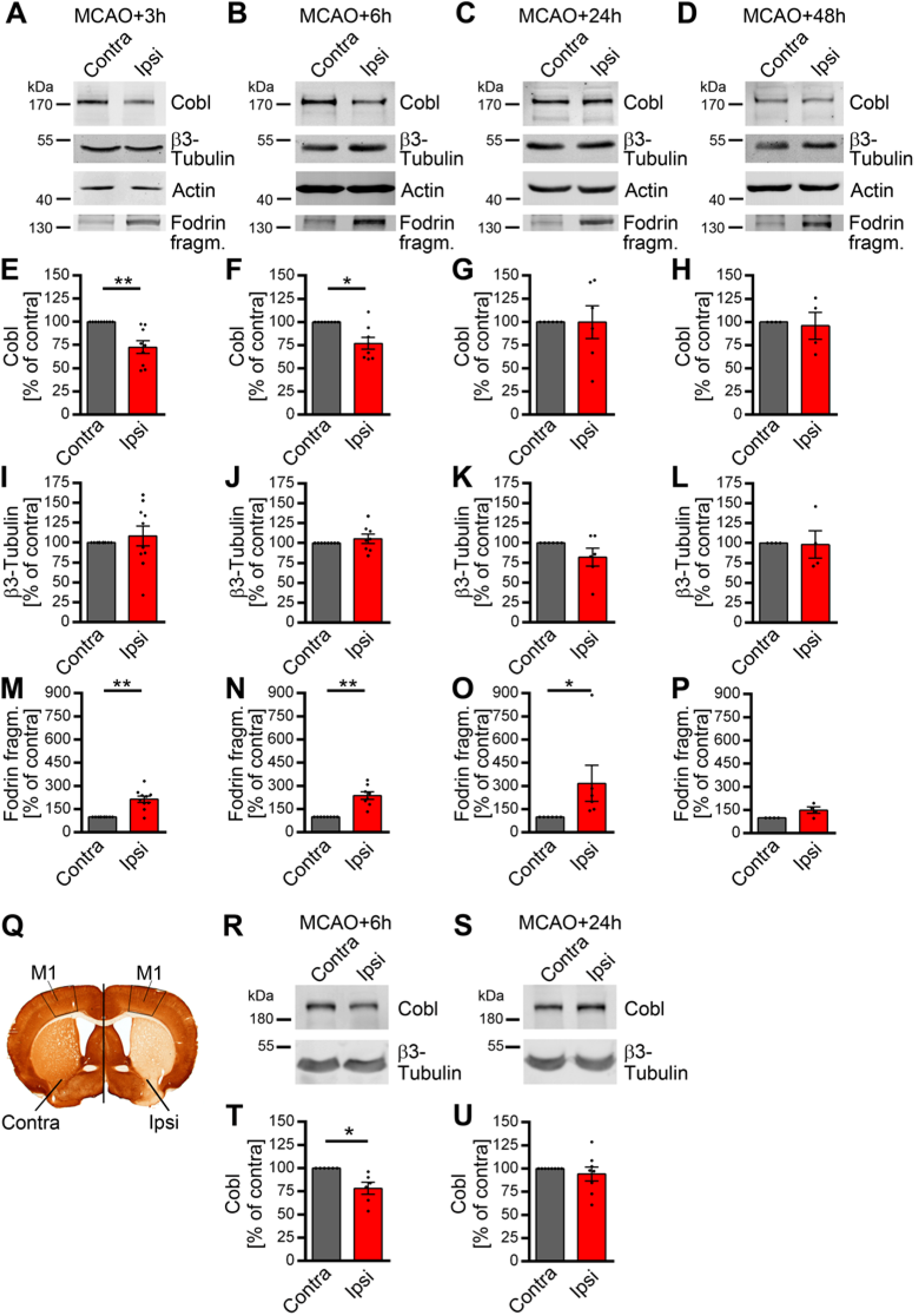
The actin nucleator Cobl decreases significantly in the ipsilateral hemisphere and in the primary motorcortex (M1) during the first hours after ischemic stroke induced by MCAO. **A-P,** Representative Western blot images (**A-D**) and quantitative analyses (**E-P**) of Cobl (**A-H**), β3-tubulin (**A-D,I-L**) and a proteolytic spectrin/fodrin fragment of 140 kDa (**A-D,M-P**) in the ipsilateral part of a middle (+0.8 and -1.2 mm to bregma) brain segment in percent of immunosignals at the corresponding contralateral side after different times of reperfusion after 30 min MCAO. Note the significant decline of Cobl protein levels at 3 h and 6 h after MCAO in the ipsilateral side (**A,B,E,F**). Anti-actin and anti-β3-tubulin immunoblotting signals served as loading controls and for normalization, respectively. **Q**, Anti-MAP2 immunostained coronal brain section with the primary motorcortex (M1) marked (black frame) at both the ipsilateral and contralateral side (dividing line) of brains of WT mice subjected to MCAO. **R-U**, Representative anti-Cobl and anti-β3-tubulin immunoblotting images (**R,S**) and quantitative analyses (**T,U**) of Cobl levels normalized to those of β3-tubulin and expressed as percent of respective control values (contra). For Cobl levels in brain hemisphere segments (+0.8 and -1.2 mm to bregma), n_MCAO+3h_=9; n_MCAO+6h_=8; n_MCAO+24h_=6; n_MCAO+48h_=4 biologically independent brain samples. For fodrin fragment and β3-tubulin levels, n_MCAO+3h_=10; n_MCAO+6h_=8; n_MCAO+24h_=6; n_MCAO+48h_=4 biologically independent brain samples. For Cobl leves in M1, n_MCAO+6h_=6; n_MCAO+24h_=8 biologically independent M1 tissue samples Data represent mean±SEM presented as bar plots overlayed with all individual data points. Statistical significance calculations, Wilcoxon signed rank test (**E-P,T,U**). \**P*<0.05; \*\**P*<0.01; \*\*\**P*<0.001.

Remarkedly, the reduced Cobl levels were rapidly restored to levels resembling contralateral values. 24 h and 48 h after MCAO, no differences in Cobl levels were observed anymore (Fig. 1C,D,G,H).

### MCAO leads to Cobl degradation followed by an efficient Cobl level recovery in the motor cortex M1

In order to directly address whether Cobl levels indeed responded to ischemic stroke conditions in wider areas of the brain, i.e. beyond the infarct area itself, and may therefore be of thus far unrecognized importance for the pathophysiology of stroke and/or subsequent repair, we conducted further MCAO experiments and isolated specifically the ipsilateral primary motor cortex (M1) - a physiologically important part of the cortex that is adjacent to but not inside the infarct area and thereby belongs to the penumbra (Fig. 1Q). Quantitative immunoblotting analyses of dissected M1 tissues clearly revealed the decline of Cobl levels at the ipsilateral side to about 75% of the corresponding contralateral control values of the respective animals 6 h after MCAO (Fig. 1R,T). Thus, the decline of Cobl in the whole brain hemisphere (Fig. 1A-H) can also clearly be observed in the penumbral M1 (Fig. 1Q-U).

Interestingly, also the recovery of Cobl expression levels at 24 h after MCAO was observed in quantitative immunoblotting analyses of M1. 24 h after MCAO, Cobl levels were restored to 100% of the control value (Fig. 1S,U).

Thus, Cobl protein levels in the cortex show surprisingly high short-term dynamics and these dynamics are related to stroke.

### Cobl degradation upon stroke is mirrored by glutamate-induced excitotoxicity brought about by NMDA receptors, high Ca^2+^ levels and calpain activity

The decreased blood flow to parts of the brain in ischemic stroke leads to a lack of oxygen and energy and therefore to a massive overrelease of glutamate and to dramatic increases of intracellular Ca^2+^ levels (Hossmann, 2006). In line with the observed degradation of Cobl in cortex samples of mice subjected to MCAO, cultures of cortical neurons subjected to increasing durations of stimulation with levels of glutamate causing excitotoxicity (Ankarcrona et al., 1995) also showed a clear and very rapid decline of Cobl protein levels in relation to unchanged actin and β3-tubulin levels (Fig. 2A,B).

**Fig. 2.**
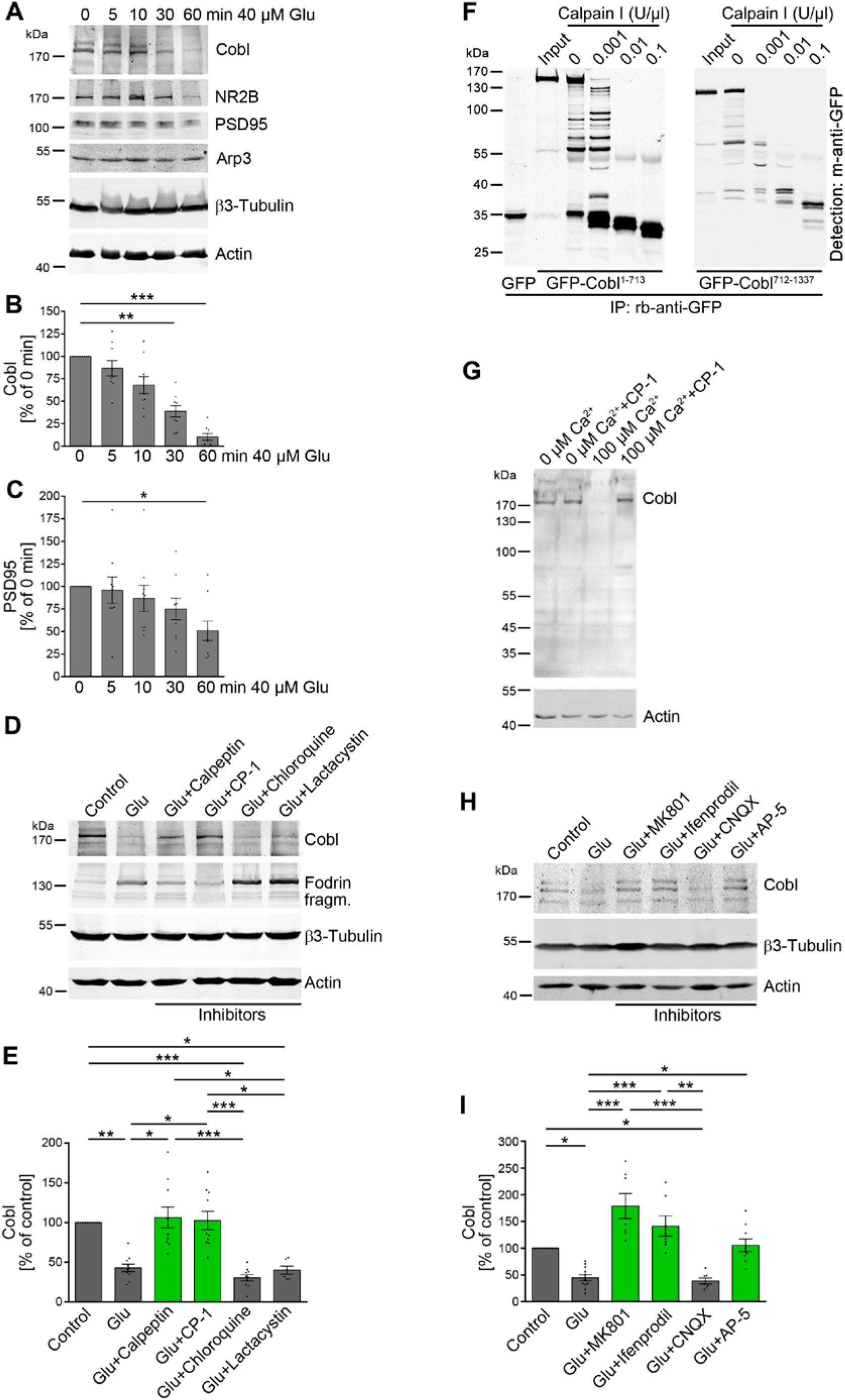
Cobl degradation upon stroke is mirrored by glutamate-induced excitotoxicity brought about by NMDA receptors, high Ca^2+^ levels and calpain activity. **A-C ,** Quantitative immunoblotting analyses of cultures of cortical neurons (DIV15) subjected to different durations of incubation with 40 µM glutamate (Glu). **A,** Representative blots. **B,C,** Excitotoxicity-induced decline of anti-Cobl (**B**) and anti-PSD95 (**C**) immunosignals. n=9 independent assays. **D,E** Proteolytic pathways underlying the glutamate-induced (30 min) Cobl decline, as shown by application of inhibitors against calpain (Calpeptin, CP-1), lysosomal degradation (Chloroquine) and proteasomal proteolysis (Lactacystin), respectively, in quantitative immunoblotting analyses. n_Control_=10, n_Glu_=10, n_Glu+Calpeptin_=10, n_Glu+CP-1_=10, n_Glu+Chloroquine_=10, n_Glu+Lactacystin_=6 biologically independent samples. **F,** Anti-GFP immunoblotting analyses of GFP-Cobl^1-713^ and GFP-Cobl^712-1337^ expressed in HEK293 cells, immunoisolated with anti-GFP antibodies (input) and incubated without (0) and with calpain I (10 min, 25°C). GFP is shown for size comparison. Note that 0.001 U/µl was sufficient for efficient Cobl digestion by calpain. **G,** Immunoblotting of brain extracts incubated (15 min, 4°C) with 0 and 100 µM Ca^2+^ and with and without the calpain inhibitor CP-1. Anti-actin signals serve as loading controls. **H,I,** NMDAR activity is required for the Cobl decline caused by excitotoxicity (40 µM glutamate, 30 min) in DIV15 cortical neurons, as shown by inhibitors against AMPA and kainate receptors (Glu+CNQX), NMDARs (Glu+AP-5), open NMDARs (Glu+MK801) and NR2B subunits of NMDA receptors (Glu+Ifenprodil), respectively, in immunoblotting (**H**) and quantitative analyses (**I**). Anti-actin and anti-β3-tubulin immunoblotting signals served as loading controls and for normalization, respectively (**A,D,H**). n_control_=12, n_Glu_=12, n_Glu+MK801_=7, n_Glu+Ifenprodil_=7, n_Glu+CNQX_=7, n_Glu+AP-5_=9 biologically independent samples. Data represent mean±SEM presented as bar plots overlayed with all individual data points. Statistical significance calculations, one-way ANOVA with Dunn’s post-test (**B,C,E,I**). \**P*<0.05; \*\**P*<0.01; \*\*\**P*<0.001. For further data also see Fig. S1.

NR2B and PSD95 are targeted by excitotoxicity processes (Simpkins et al., 2003; Gascón et al., 2008). The Cobl decline was more rapid and more severe than the declines observed for the established excitotoxicity-induced degradation targets NR2B and PSD95 (Fig. 2A-C; Fig. S1A).

In contrast to the actin nucleator Cobl and to the postsynaptic components NR2B and PSD95, quantitative anti-Arp3 evaluations did not show any declines in Arp2/3 complex levels (Fig. 2A; Fig. S1B). This was somewhat surprising as MCAO is known to modulate dendritic spines (Brown et al., 2008; Wu et al., 2014) and the Arp2/3 complex and modulators of its activity, respectively, have been thoroughly established as important for postsynaptic F-actin organisation shaping dendritic spines in multiple ways (Kim et al., 2006; Soderling et al., 2007; Haeckel et al., 2008; Wegner et al., 2008; Spence et al., 2016). On the other hand, only very few proteins, such as NR2B and PSD95 (Simpkins et al., 2003; Gascón et al., 2008), have thus far been identified as directly responsive to excitotoxicity. Arp3 may be protected from proteolytic degradation by its compact fold and its tight interactions with the other Arp2/3 complex components as well as with actin.

Cobl protein levels declined upon excitotoxicity paradigms in both cultures and animals. Excitotoxicity processes are linked to high intracellular Ca^2+^ levels. Consistently, the application of different protease inhibitors showed that the Cobl decline was brought about by the Ca^2+^-activatable protease calpain (Fig. 2D,E; Fig. S1C). Both calpain inhibitors, calpain inhibitor 1 (CP-1) and Calpeptin, did not only suppress the occurrence of spectrin/fodrin fragments upon glutamate application but also effectively inhibited the Cobl decline (Fig. 2D,E). Chloroquine and Lactacystin, in contrast, did not suppress the excitotoxicity-induced decline of Cobl protein levels (Fig. 2D,E). Thus, the decline of Cobl did neither reflect a lysosomal nor a proteasomal degradation but was calpain-mediated.

Reconstitutions with immunoprecipitated Cobl N and C terminal halves, respectively, and rising calpain concentrations clearly demonstrated that calpain can indeed lead to a rapid Cobl destruction (Fig. 2F). A further mapping of calpain cleavage sites revealed that even the use of smaller parts of Cobl (GFP-Cobl^1-408^, GFP-Cobl^406-866^, GFP-Cobl^750-1005^ and GFP-Cobl^1001-1337^) still showed effective degradations into multiple, GFP-containing fragments (Fig. S1D). The multitude of calpain cleavage sites suggest a complete loss of Cobl functions when Cobl is digested by calpain.

Also in brain extracts, Cobl was susceptible to Ca^2+^-triggered degradation. Importantly, this degradation of endogenous Cobl again was suppressible by calpain inhibitor (Fig. 2G).

We next addressed the neuronal signal transduction pathways leading to the calpain-mediated Cobl digestion. CNQX, a powerful inhibitor of AMPA and Kainate types of glutamate receptors, did not suppress the excitotoxicity-mediated Cobl decline. In contrast, AP-5, MK801 and also Ifenprodil, which generally interfere with NMDARs, inhibit open NMDARs and explicitly block the NR2B subunit of NMDARs, respectively, all completely suppressed Cobl degradation caused by prolonged incubation with 40 µM glutamate (Fig. 2H,I; Fig. S1E for β3-tubulin). Thus, the glutamate-induced Cobl degradation is mediated by the activation and opening of NR2B-containing NMDARs.

Serial section analyses of brains of *Cobl* KO mice (Haag et al., 2018) subjected to 30 min MCAO showed that neither the ischemia-induced acute loss of Cobl we detected in the central brain segment and in the M1 nor the subsequent rapid restoration of normal Cobl levels in these brain parts had any influence on the survival of neurons within the infarct area in the striatum (Fig. S1F-H).

With about 5% of the whole brain affected, *Cobl* KO mice had infarct volumes comparable to those of WT mice (Fig. S1F-H). This extend of lesional damage caused by 30 min MCAO in both genotypes also reliably mirrored the range of survivable human stroke, which usually also represents about 5% of the total brain (Cramer et al., 2006).

### A transient increase of *Cobl* mRNA preceeds the rapid recovery of Cobl protein levels after its post-stroke decline

The observed rapid recovery of ipsilateral Cobl protein levels 24 h after MCAO (Fig. 1) suggested active counteractions of the stroke-affected brain to rapidly replenish the actin nucleator Cobl for some important, yet unknown function(s). This compensation mechanism should be reflected by increased mRNA levels at specifically the ipsilateral, i.e. the stroke-affected side, prior to full restoration of normal Cobl protein levels at 24 h after MCAO. We indeed observed a statistically highly significant increase of *Cobl* mRNA levels at the ipsilateral side at two different time points prior to restoration of Cobl protein levels to contralateral control levels at 24 h after MCAO (Fig. 3A).

**Fig. 3.**
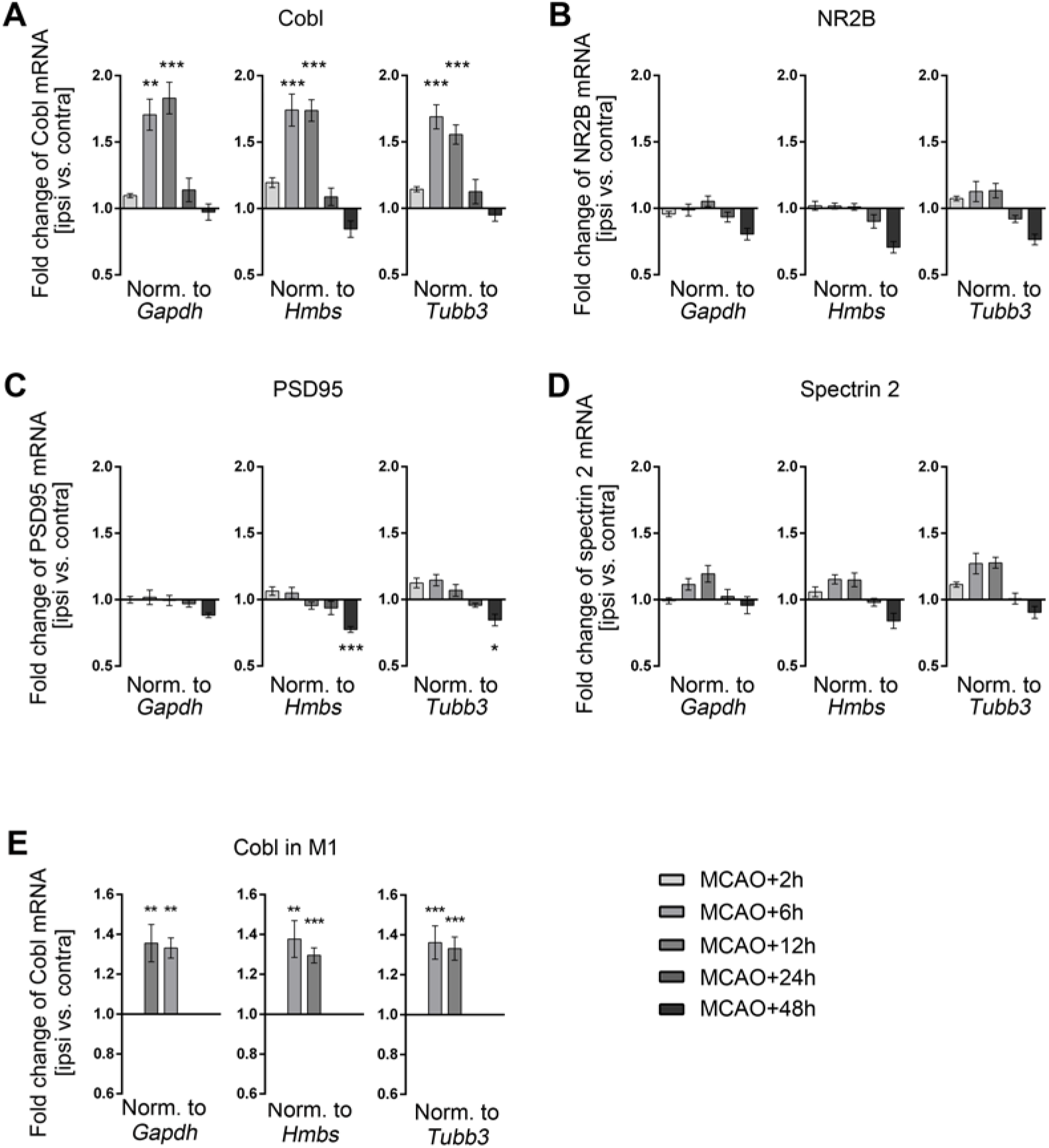
qPCR analyses demonstrate a transient increase of *Cobl* mRNA levels following the decline of Cobl protein levels upon MCAO. **A-D ,** Fold change of mRNA levels of *Cobl* (**A**), *NR2B* (**B**), the scaffolding proteins *PSD95* (**C**) and *spectrin 2* (**D**) determined in the middle segment of the brain (+0.8 and -1.2 mm to bregma) by qPCR after 2 h, 6 h, 12 h, 24 h and 48 h of reperfusion after 30 min MCAO. Data represent the differences of *Cobl*, *NR2B*, *PSD95* and *spectrin 2* mRNA levels (ipsi vs. contra) normalized to *Gapdh* (left panel), to *Hmbs* (middle panel) and to the neuronally expressed gene *Tubb3* (right panel), respectively. Brain samples from 6-9 mice were analyzed for each time point. **E,** Fold changes of *Cobl* mRNA levels in M1 tissue samples (ipsi vs. contra) at the times of upregulation identified above (6 h and 12 h after MCAO). The data was again normalized against three different genes (*Gapdh*, *Hmbs* and *Tubb3*). **A,** n_MCAO+2h_=8; n_MCAO+6h_=8; n_MCAO+12h_=9; n_MCAO+24h_=6; n_MCAO+48h_=6 mice. **B-D,** n_MCAO+2h_=7; n_MCAO+6h_=8; n_MCAO+12h_=7; n_MCAO+24h_=6; n_MCAO+48h_=6 mice. **E,** n_MCAO+6h_=7; n_MCAO+12h_=8. Data represent mean±SEM. Statistical significances (ipsi vs. contra) were calculated using one-way ANOVA with Sidak’s post-test. *\*P*<0.05; *\*\*P*< 0.01; *\*\*\*P*< 0.001. For further data also see Fig. S2.

The increase of *Cobl* mRNA levels in the ipsilateral hemisphere of the brain was detected as early as 6 h after MCAO (Fig. 3A), i.e. during a time when Cobl protein levels still were strongly negatively affected (Fig. 1B,F). The increase of *Cobl* mRNA 6 h after MCAO was validated using two additional primer pairs. The results were fully consistent irrespective of primers and normalizations used (Fig. 3A; Fig. S2A,B). The increase of *Cobl* mRNA levels after MCAO remained equally strong at 12 h (Fig. 3A). It then ceased 24 h after MCAO (Fig. 3A) when also Cobl protein levels reached a level similar to the contralateral control side (Fig. 1C,G).

The increase of *Cobl* mRNA levels reflected changes in especially neurons, as it occurred upon normalization to *Gapdh* and *Hmbs* mRNA levels (ubiquitously expressed house-keeping genes) but also upon normalization to *Tubb3* mRNA levels (Fig. 3A; Fig. S2A,B).

Interestingly, although NR2B, PSD95 and spectrin 2/fodrin also are targets of proteolytic processes triggered by excitotoxicity (Simpkins et al., 2003; Pike et al., 2004; Gascón et al., 2008), the mRNA of none of these components showed any transient increase of expression levels after MCAO (Fig. 3B-D). The levels of *CaM 1* and *calpain 1* mRNA - i.e. of two factors that dictate the activity and the functionality, respectively, of the actin nucleator Cobl - also did not change at any examined time point subsequent to MCAO (Fig. S2C,D). The transient increase of *Cobl* mRNA levels subsequent to the exitotoxicity-mediated Cobl digestion by calpain and in advance to the rebound of Cobl protein levels thus seems to be a unique compensatory response to ischemic stroke.

This raised the question whether this stroke response could indeed be observed in penumbral areas, such as the M1 similar to the changes in Cobl protein levels (Fig. 1). We therefore expanded our Cobl analyses in M1 tissue by qPCR experiments at both time points of elevated *Cobl* mRNA levels using two independent primer pairs (Fig. 3E; Fig. S2E). Cobl mRNA showed clear upregulations in both 6 h and 12 h post-MCAO tissue samples of M1 (Fig. 3E, Fig. S2E).

Thus, both the Ca^2+^-, calpain- and NMDA receptor-mediated decline of Cobl as well as its fast recovery to normal levels driven by a transient increase in *Cobl* mRNA were phenomena observable in neurons of a penumbral cortex area, the M1.

### MCAO leads to strong reductions of dendritic arbor complexity in layers II/III and V of the primary motor cortex (M1)

Besides axons with their presynapses and dendritic spines harboring postsynapses, also the dendritic arbor itself is a major structural element in neuronal wiring in the brain. As the formation of new synaptic contacts subsequent to stroke-dependent loss (Brown et al., 2008; Wu et al., 2014) may only be one aspect in the penumbra that compensates for the functions of entire brain regions lost upon ischemic stroke and recently also acute penumbral dynamics of the dendritic arbor were reported after ischemic stroke (Hu et al., 2019; Mauceri et al., 2020), we analyzed the dendritic arbor of the ipsilateral M1.

In order to obtain reliable data addressing infarct-related dendritic changes, we established a procedure that enabled us to evaluate the core infarct area caused by 30 min MCAO in the striatum of each individual mouse by anti-MAP2 immunostaining and to in parallel analyze morphologies of individual neurons in a defined area adjacent to the damage zone, the penumbra (represented by the primary motor cortex (M1)) using neighbored coronal brain sections (Fig. S3A,B).

Detailed morphometric analyses of both layer II/III (Fig. 4A-C) and layer V neurons of the ipsilesional M1 (Fig. 4D-F) unveiled that MCAO leads to dramatic dendritic arborization defects when compared to sham-treated animals, which also underwent anesthesia and the surgical procedures but without MCAO (Fig. 4A-N). Both the number of dendritic branching points and the number of terminal points per neuron declined subsequent to MCAO at specifically the ipsilateral side when brains were analyzed after 24 h reperfusion (Fig. 4G,H,K,L).

**Fig. 4.**
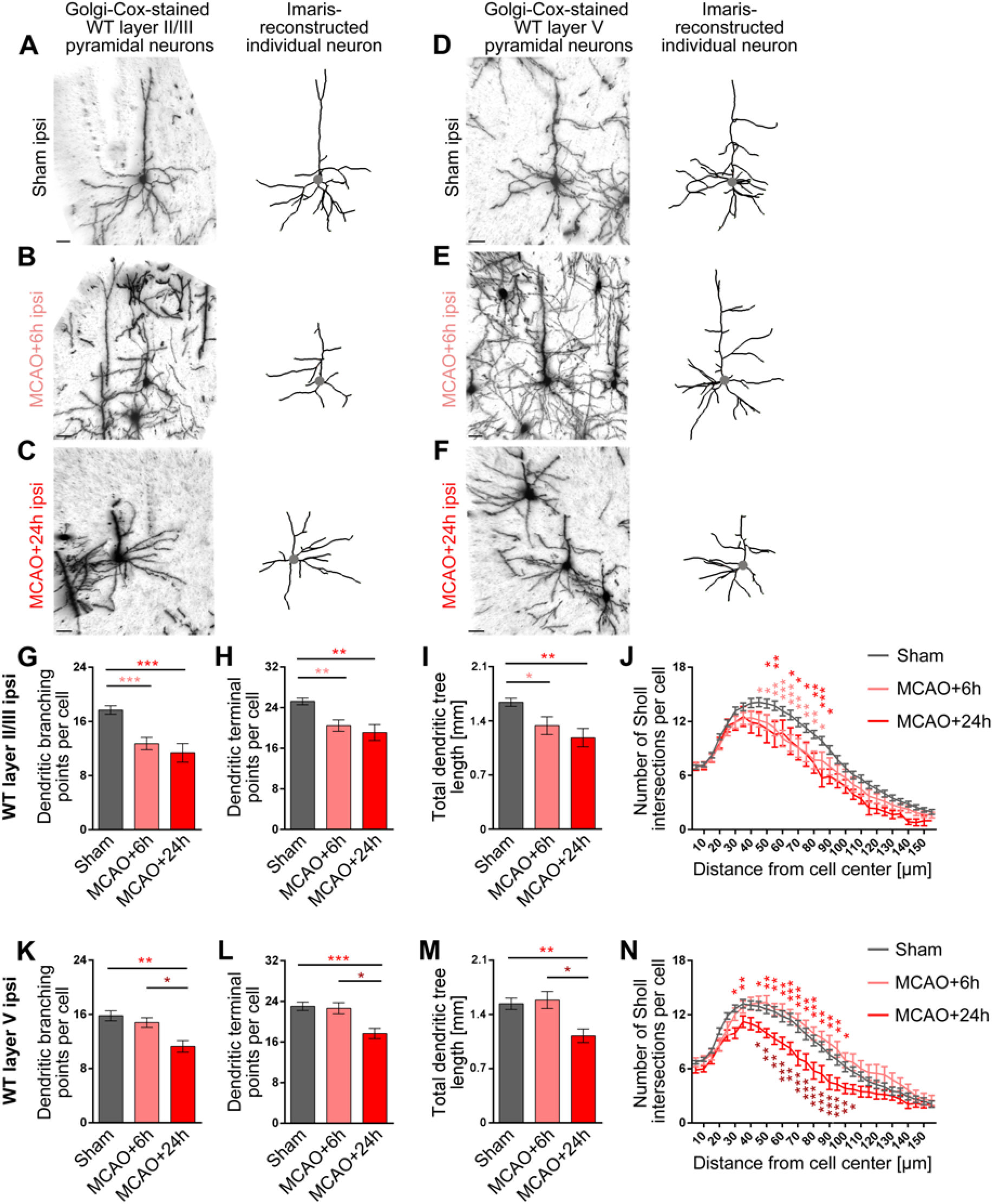
Loss of dendritic arbor complexity subsequent to ischemic stroke in ipsilateral layer II/III and layer V neurons of the motor cortex. **A-F,** Representative images of Golgi-Cox-stained (left panels) and individual Imaris-reconstructed (right panels) layer II/III (**A-C**) and layer V (**D-F**) pyramidal neurons of M1 from the ipsilateral side of sham-treated control mice (**A,D**) and of mice subjected to 30 min MCAO analyzed after 6 h (**B,E**) and 24 h (**C,F**) reperfusion times. The position of the cell bodies are marked by a gray dot. Scale bars, 30 μm. **G-N,** Quantitative determinations of dendritic branching points, terminal points, total dendritic tree length and Sholl intersections of dendritic trees in layer II/III (**G-J**) and layer V (**K-N**). Note that all parameters of dendritic complexity at the ipsilateral side decreased subsequent to MCAO and that layer II/III neurons responded faster, as they already show the decreased dendritic arborization at 6 h after MCAO (for contralateral data see Fig. S3). Layer II/III: n_Sham_=60; n_MCAO+6h_=22; n_MCAO+24h_=11 neurons. Layer V: n_Sham_=47; n_MCAO+6h_=19; n_MCAO+24h_=18 neurons from 3 mice for each MCAO group and 6 mice for the sham control (ipsi). Quantitative data represent mean±SEM. Statistical significance calculations, one-way ANOVA with Tukey’s post-test (**G-I,K-M**) and two-way ANOVA with Sidak’s post-test for Sholl analysis (**J,N**), respectively. *\*P<0.05*; *\*\*P<0.01*; *\*\*\*P<0.001*.

In cortex layer II/III, neurons even showed these dramatic impairments already after 6 h (Fig. 4G,H). The MCAO-mediated defects in layer V neurons developed slower and were observable at 24 h (Fig. 4K,L). Similar defects and onsets were observed when the entire length of the dendritic arbor was examined in layer II/III and layer V of the cortex (Fig. 4I,M). Also Sholl analyses of the dendritic complexity showed corresponding reductions of the dendritic arbor (Fig. 4J,N).

These ischemic stroke-induced defects in dendritic organization of neurons in both analyzed areas of the motor cortex were restricted to the ipsilesional side. For all four parameters determined the data obtained from the contralateral side of the same MCAO animals were indistinguishable from those of sham-treated mice (Fig. S3C-M).

### As early as 4 days after MCAO, the discovered dendritic arborization defects caused by MCAO were mostly compensated

We next evaluated whether the massive MCAO-induced impairment in dendritic arborization we observed in the M1 (Fig. 4) was permanent and part of the lesion-based disabilities caused by stroke or whether it would at some point become repaired by some thus far unidentified form of dendritic dynamics, which may be inducible by ischemic stroke in mature neurons. We therefore analyzed the morphologies of pyramidal neurons in the M1 of WT mice at 4 days and 7 days after MCAO in direct comparison to the defects observed at 24 h and to sham controls in a blinded manner (Fig. 5A-D). Strikingly, in neurons of the ipsilateral layer II/III, all defects observed at 24 h after MCAO were fully compensated for at day 4 and 7 (Fig. 5E-J). The dendritic branching points at day 4 and 7 after MCAO exactly were at the levels of sham animals (Fig. 5E). Likewise, dendritic terminal points were back at control levels (Fig. 5F). Total dendritic tree length and Sholl analyses also showed a full restoration of normal dendritic arborization at 4 days and 7 days after MCAO (Fig. 5G-J).

**Fig. 5.**
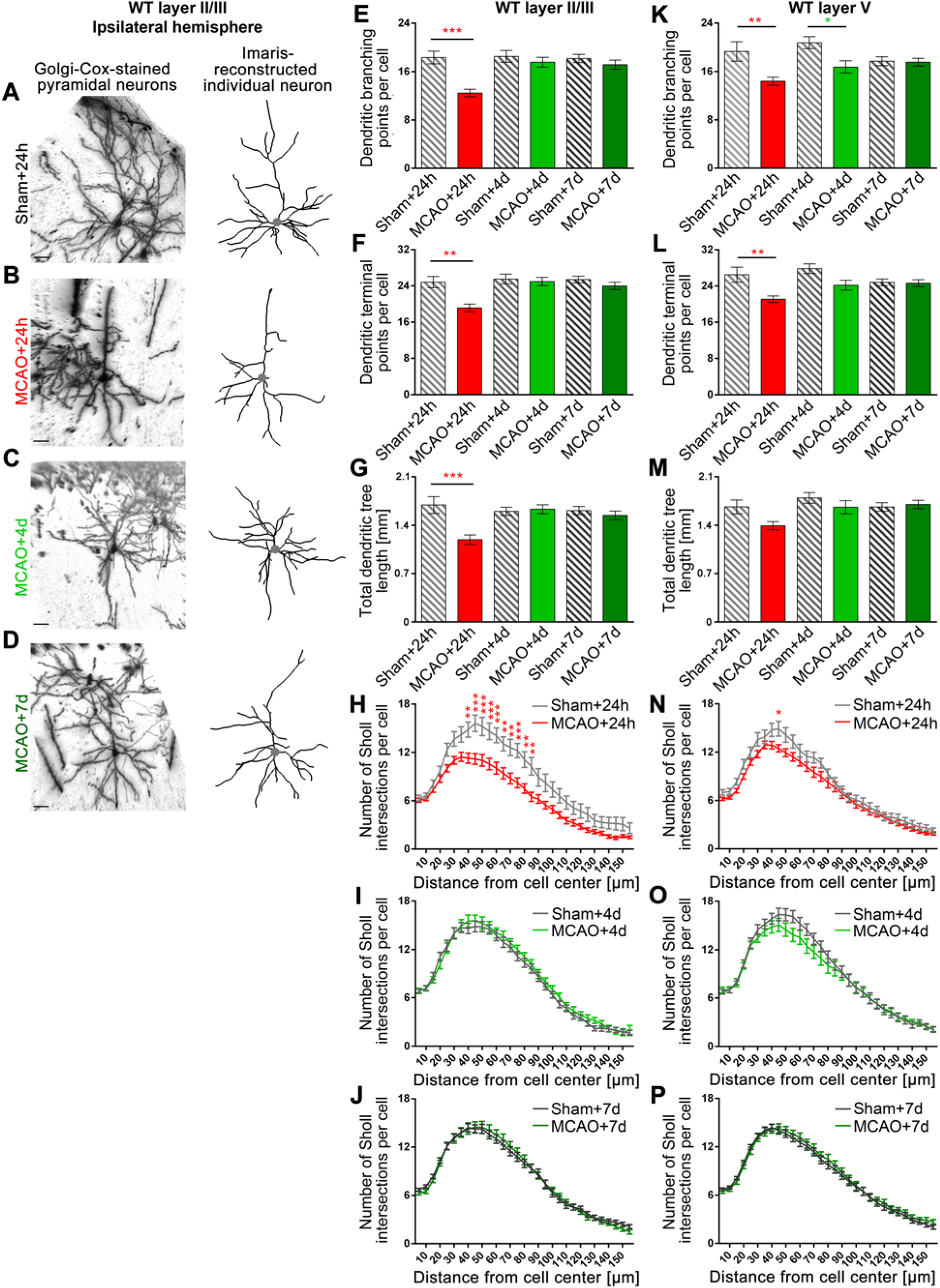
Dendritic arborization defects caused by MCAO in M1 are completely reverted during the days after the ischemic event in both layer II/III and layer V neurons of the ipsilateral hemisphere. **A-D,** Representative images of Golgi-Cox-stained and Imaris-reconstructed ipsilateral layer II/III pyramidal neurons of M1 of WT mice subjected to sham surgery and of WT mice subjected to 30 min MCAO at 24 h and 4 d, 7 d of reperfusion time. The position of the cell bodies are marked by a gray dot. Scale bars, 30 μm. **E-J,** Quantitative determinations of dendritic arborization parameters of layer II/III neurons. Note that dendritic branching points (**E**), dendritic terminal points (**F**), total dendritic tree length (**G**) and Sholl intersections (**H-J**) all were decreased upon MCAO after 24 h but were fully recovered to values indistinguishable from the corresponding sham controls after 4 d and 7 d reperfusion, respectively. **K-P,** Quantitative analyses of dendritic arborization parameters of layer V neurons at 24 h, 4 d and 7 d reperfusion. Ipsilateral M1 tissue of 3 mice of an age of 3-4 months were analyzed for the sham+24h, sham+4d and MCAO+4d groups and 6 mice for the sham+7d, MCAO+24h and MCAO+7d groups, as two independent animal cohorts were evaluated and included (for initial MCAO+24h data from only 3 mice e.g. see Fig. 4) (for contralateral data see Fig. S5). Layer II/III: n_Sham+24h_=18; n_MCAO+24h_ =43; n_Sham+4d_=35; n_MCAO+4d_=32; n_Sham+7d_=63; n_MCAO+7d_=47 neurons. Layer V: n_Sham+24h_=22; n_MCAO+24h_=51; n_Sham+4d_=27; n_MCAO+4d_=29; n_Sham+7d_=61; n_MCAO+7d_=51 neurons. Data represent mean±SEM. Statistical significances calculated between sham and MCAO of the corresponding reperfusion time using two-way ANOVA with Sidak’s post-test are shown. *\*P<0.05*; *\*\*P<0.01*; *\*\*\*P<0.001*.

Both apical and basal parts of the dendritic arbor of layer II/III neurons showed about equally strong defects (Fig. S4A-K). Dendritic branch points and total dendritic length were reduced by about 30% (Fig. S4D,F,I,K). In both apical and basal dendrites, the terminal point numbers declined by 25-30% but failed to reach statisitcal significance for the apical dendrites (Fig. S4E,J). Also the restoration of dendritic arbor complexity was as effective in apical dendrites as it was in basal dendrites (Fig. S4D-K).

Layer V neurons, which showed a slower onset of dendritic defects (Fig. 4), showed a similarly clear repair of MCAO-induced impairments (Fig. 5K-P). Only the number of dendritic branching points still showed some reduction at day 4 (Fig. 5K). At day 7 after MCAO, all dendritic morphology parameters quantitatively analyzed in MCAO-treated animals were no longer significantly different from those obtained from sham animals (Fig. 5K-M,P).

Together, our evaluations at the different time points unveiled that the dendritic arborization defects, which we detected upon MCAO in both layer II/III and layer V of the M1, were fully compensated for by a dendritic regrowth process, which gave rise to apparently normally branched and sized dendritic trees and which occured in both layer II/III and layer V.

Examinations at the contralateral side showed that the ischemic stroke-triggered repair process in M1 is restricted to the defective ipsilateral side and is not a brain-wide phenomenon of dendritic growth induction. At the contralateral side, all parameters remained at sham levels during all time points analyzed (Fig. S5).

Severe ischemic stroke models, such as 60 min MCAO or even a permanent blockage of blood flow in mice, being rather unrelated to survivable human strokes, would not be suitable to study repair processes in the penumbra, as surving animals usually lose large parts of the entire hemisphere affected and show significant structural changes at distant sites or even at the contralateral side of the brain (Cramer et al., 2006; Winship and Murphy, 2008) and the penumbra is small or not existing (Popp et al., 2009). We did not observe any structural alterations at the contralateral side (Fig. S5) but specifically found cellular alterations in the penumbra at the ipsilateral side (Figs 4 **and** 5). This demonstrated that our animal stroke model (30 min MCAO) reliably mirrored the range of survivable human stroke and thus was suitable to unveil repair processes in peri-infarct areas.

### The actin nucleator Cobl promotes and is critical for dendritic arborization in cortical neurons

The defects caused by MCAO in pyramidal neurons in cortical layers II/III and V of the murine primary motor cortex seemed somewhat related to those observed for Cobl loss-of-function in immature rat hippocampal neurons in culture (Ahuja et al., 2007; Izadi et al., 2021). We therefore addressed whether Cobl may also play some important role in shaping cortical neurons and whether furthermore Cobl may not do so not only in developing neurons in culture but also in adult neurons in the cortex of mice with a demand for remapping neuronal circuits subsequent to stroke.

Adressing the first hypothesis, we overexpressed Cobl in developing rat cortical neurons. GFP-Cobl overexpression from DIV4 to DIV6 clearly resulted in increased dendritic arborization when compared to GFP control. All parameters affected upon MCAO in mice were elevated significantly (Fig. S6A-F).

In line with these results, Cobl loss-of-function experiments in developing primary rat cortical neurons unveiled that Cobl does not only have the ability to modulate the dendritic trees of cortical neurons but, but also is critical for this process during dendritic arbor developement (Fig. 6A-C). Dendritic branching points, terminal points and the total dendritic length were significantly reduced and also Sholl analyses highlighted a loss of dendritic complexity when Cobl was lacking (Fig. 6D-G). All of these Cobl loss-of-function phenotypes were specific, as all of them could be rescued by reexpression of RNAi-insensitive Cobl in the cultured cortical neurons (Fig. 6C-G).

**Fig. 6.**
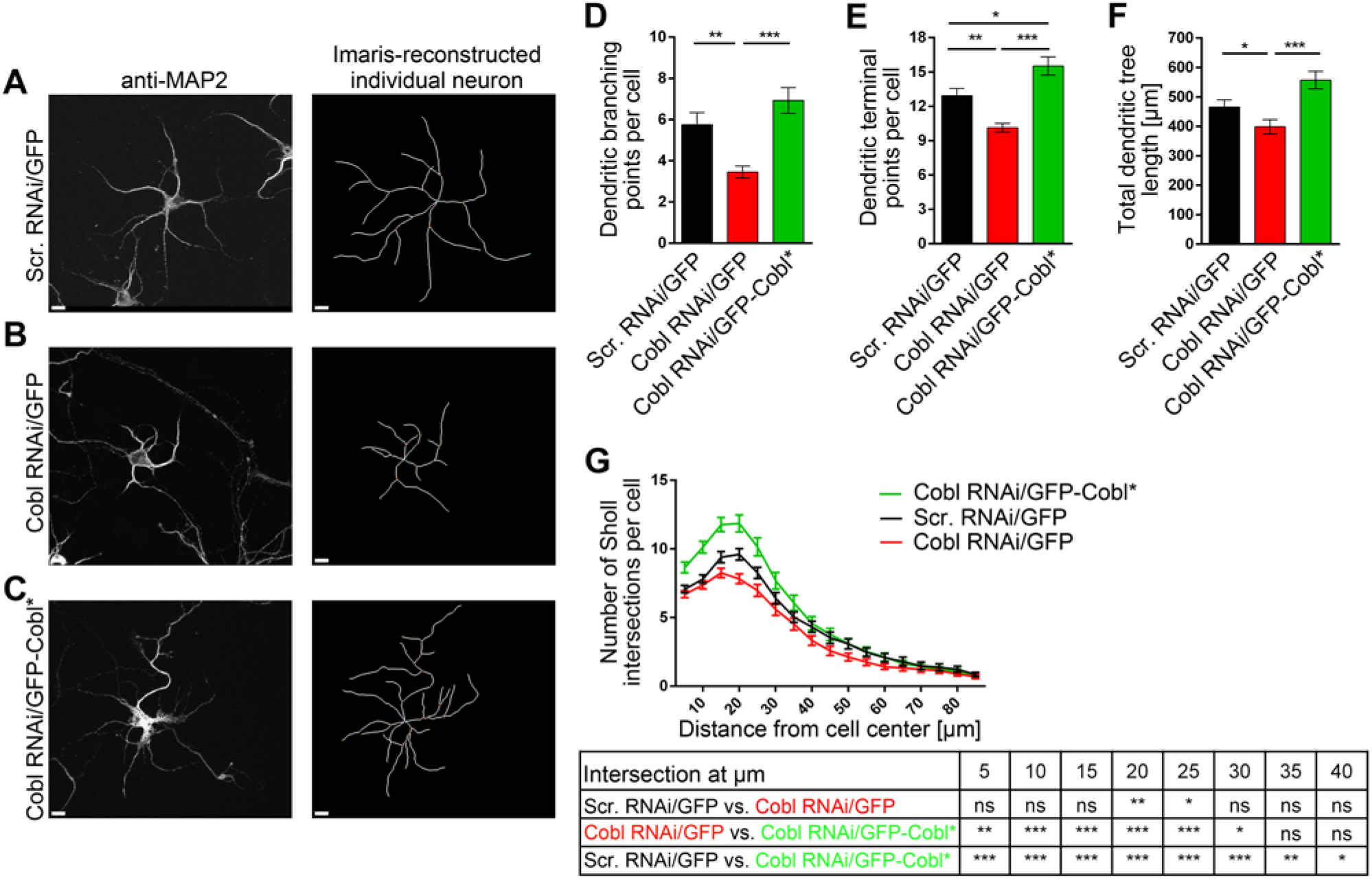
The actin nucleator Cobl is critical for dendritic arborization during early development of cultured primary rat cortical neurons. **A-C ,** Representative maximum intensity projections (MIPs) of anti-MAP2-immunostained (left panels) and Imaris-reconstructed primary rat cortical neurons (right panels) that were transfected at DIV4 with plasmids encoding for either GFP-reported scrambled RNAi (Scr. RNAi/GFP) (**A**), Cobl RNAi/GFP (**B**) and Cobl RNAi together with silently mutated Cobl rendered insensitive again Cobl RNAi (Cobl RNAi/GFP-Cobl*) (**C**), respectively, and fixed 40 h later. Scale bars, 10 μm. **D-G,** Quantitative determinations of dendritic arborization parameters unveiling clear and specific Cobl loss-of-function phenotypes. Note that dendritic branching points (**D**), dendritic terminal points (**E**), total dendritic tree length (**F**) and Sholl intersections (**G**) all were decreased upon Cobl RNAi and were rescued to levels clearly statistically different from Cobl RNAi by reexpressing GFP-Cobl*. n_Scr. RNAi/GFP_=40; n_Cobl RNAi/GFP_=40; n_Cobl RNAi/GFP-Cobl*_=40 individual transfected neurons from 2 independent preparations of cortical neurons. Data, mean±SEM. Statistical significances were calculated using One-way ANOVA with Tukey post-test (**D-F**) and two-way ANOVA with Sidak’s post-test for Sholl analysis (**G**). *\*P*< 0.05; *\*\*P*< 0.01; *\*\*\*P*< 0.001.

Taken together, these two different experimental lines clearly proved the first hypothesis and unveiled that the actin nucleator Cobl played an important role in the dendritic arborization of cortical neurons.

### *Cobl* KO completely ablates the ischemic stroke-induced dendritic regrowth processes, which allow for restoration of proper dendritic arborization after MCAO

The second hypothesis, that, as reflected by the identified modulations of Cobl levels acutely following ischemia and during stroke recovery, the actin nucleator Cobl may be a crucial player in the repair of the stroke-induced dendritic arbor defects in the penumbra within the observed narrow time window between day 1 and day 4 after MCAO, was addressed by subjecting *Cobl* KO mice (Haag et al., 2018) to comparative and time-resolved MCAO studies evaluating the different dendritic parameters in a fully blinded manner. Similar to WT mice (Fig. 5), also *Cobl* KO mice showed a reduction of dendritic branching points, terminal points and total dendritic tree length 24 h after MCAO. At 24 h after MCAO, the reduction in each parameter was about 25% in *Cobl* KO mice (Fig. 7E-G,K-M). These MCAO-induced declines were about as strong as detected in WT animals (Fig. 5). Deviations in Sholl analyses of dendritic arbor complexity of *Cobl* KO neurons very well reflected the reductions in dendritic branching points, terminal points and tree extension (Fig. 7H,N). In general, all individual defects in *Cobl* KO mice were about equally strong in layer II/III and in layer V 24 h after MCAO (Fig. 7E-G,K-M).

**Fig. 7.**
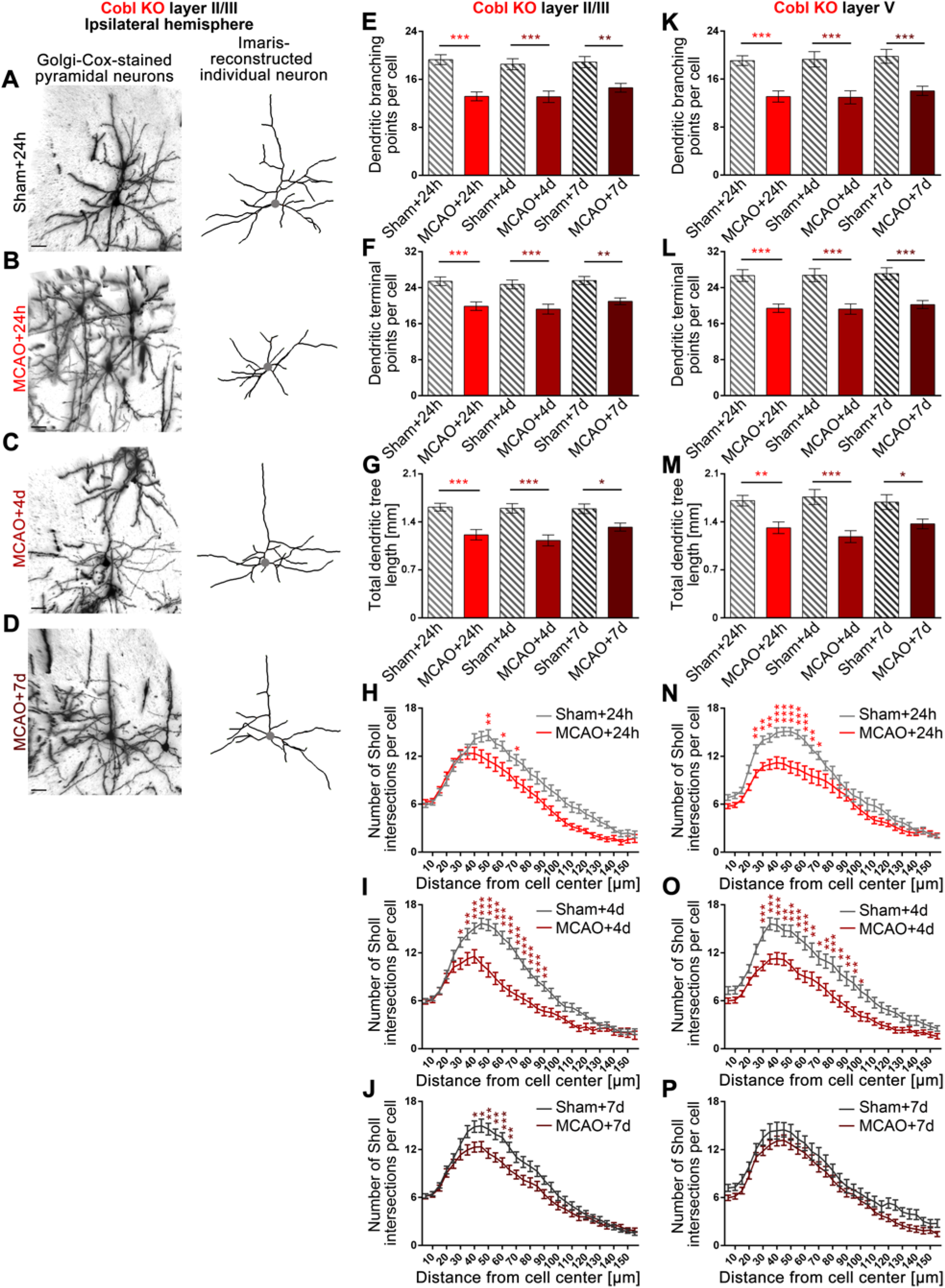
*Cobl* KO mice fail to show repair of the dendritic arborization defects caused by MCAO in both layer II/III and layer V neurons of the ipsilateral hemisphere. **A-D ,** Representative images of Golgi-Cox-stained and Imaris-reconstructed ipsilateral layer II/III pyramidal neurons of M1 of *Cobl* KO mice subjected to sham surgery and of *Cobl* KO mice subjected to 30 min MCAO at 24 h, 4 d and 7 d of reperfusion time. The position of the cell bodies are marked by a gray dot. Scale bars, 30 μm. **E-J,** Quantitative determinations of dendritic arborization parameters of layer II/III neurons. Note that dendritic branching points (**E**), dendritic terminal points (**F**), total dendritic tree length (**G**) and Sholl intersections (**H-J**) all were decreased at 24 h after MCAO and that, in contrast to WT animals (Figure 5), none of these phenotypical parameters recovered during the days after MCAO in *Cobl* KO mice. Instead, all dendritic parameters remained strongly suppressed even after 4 d and 7 d of reperfusion. **K-P,** Quantitative analyses of dendritic arborization parameters of layer V neurons in M1 at 24 h, 4 d and 7 d reperfusion showing a similar lack of dendritic recovery from ischemic stroke upon *Cobl* KO. For each MCAO and sham group, 3 mice of an age of 3-4 months were analyzed (for contralateral data see Fig. S7). Layer II/III: n_Sham+24h_=21; n_MCAO+24h_=35; n_Sham+4d_=30; n_MCAO+4d_=29; n_Sham+7d_=26; n_MCAO+7d_=35 neurons. Layer V: n_Sham+24h_=25, n_MCAO+24h_=31, n_Sham+4d_=21, n_MCAO+4d_=25, n_Sham+7d_=19, n_MCAO+7d_=30 neurons. Data represent mean±SEM. Statistical significances calculated between sham and MCAO of the corresponding reperfusion time using two-way ANOVA with Sidak’s post-test are shown. *\*P<0.05; **P<0.01; ***P<0.001*.

Again similar to WT mice, also *Cobl* KO mice did not show any significant MCAO-induced neuronal morphology changes on the contralateral side (Fig. S7) but MCAO-induced changes were restricted to the ipsilateral side (Fig. 7).

By evaluating the restoration of proper dendritic arbor complexity after MCAO at the ipsilateral side, it became obvious that the repair of dendritic arborization subsequent to MCAO was grossly impaired in *Cobl* KO mice. All four quantified dendritic defects remained fully present 4 days after MCAO. Neither dendritic branching points (Fig. 7E,K), nor dendritic terminal points (Fig. 7F,L) nor dendritic tree length (Fig. 7G,M) nor dendritic complexity, as visualized by Sholl analyses (Fig. 7I,O), were restored to proper levels in *Cobl* KO animals. This phenotype became obvious by comparision to sham-treated animals (Fig. 7) but also by comparing the absolute numbers of all parameters with those of WT animals (Fig. 7 vs. Fig. 5).

At day 4, this complete lack of repair of MCAO-induced defects was observed in both layer II/III neurons (Fig. 7C,E-G,I) and in layer V neurons of *Cobl* KO mice (Fig. 7K-M,O). The lack of repair at day 4 also occurred irrespective of the dendritic orientation, as MCAO-induced defects in *Cobl* KO neurons were observed in both the apical and the basal dendritic arbor (Fig. S8).

Even at day 7 after MCAO and again in strong contrast to WT animals, *Cobl* KO mice still showed massive impairments in dendritic arborization (Fig. 7D,E-G,J,K-M). Only Sholl analyses of layer V neurons failed to visualize the differences in dendritic complexity between sham- and MCAO-treated *Cobl* KO mice in a manner supported by statistical significances (Fig. 7P). Yet, all other seven quantitative examinations still clearly demonstrated the lack of any dendritic arbor restoration in *Cobl* KO mice even a full week after MCAO (Fig. 7E-P). The dendritic branching points, the terminal points and the summarized length of the dendritic arbor remained significantly reduced when compared to sham-treated animals at day 7. The defects at day 7 thereby still remained similar to the defects observed at 24 h and 4 d (Fig. 7E-P). Taken together, the levels of the actin nucleator Cobl are responsive to MCAO-induced brain damage and Cobl is critically needed for an acute process of dendritic regrowth and branching that is triggered by ischemic stroke conditions and is powerful enough to fully repair the strong defects in dendritic arborization, which we observed as consequences of ischemic stroke in the cortex.

## Discussion

Regaining of neurophysiological functions in peri-infarct areas (the penumbra) and remapping processes in functionally related cortical tissues are thought to represent important aspects underlying recovery from stroke in human patients. However, such processes are far from being understood at the cellular and molecular level.

We demonstrated that ischemic stroke in mice leads to a strong decline in the entire dendritic arborization of neurons in the penumbra and by carefully comparing different reperfusion times with each other and sets of corresponding controls demonstrate a subsequent process of dendritic arbor repair. In the cortex of mice subjected to 30 min MCAO-induced ischemic stroke in the striatum, this repair process showed a net growth that was able to fully compensate for the massive loss of dendritic extension and complexity observed at the first day after ischemic stroke. Importantly, with the actin nucleator Cobl, we identified a cytoskeletal driving force for this recovery. Dendritic arbor repair was completely absent in *Cobl* KO mice subjected to ischemic stroke. Neurons in M1 of *Cobl* KO mice subjected to MCAO remained at the state of severe dendritic impairment.

During the repair in WT animals, the observed growth represented an increase of the dendritic arbor by one third above 24 hours post-stroke levels. Such significant changes in dendritic arborization in adult animals are quite stunning, as the dendritic arbor, in contrast to dendritic spines, which are known to at least retain some minor dynamics in adults, is established to be rather resilient under normal conditions (Koleske, 2013). Recent observations in rats subjected to transient MCAO (Hu et al., 2020), however, are fully in line with the drastic changes we observed in mice. Although not fully comparable to our studies, as a stonger MCAO was used (90 min), ischemic stroke was reported to result in significant declines in the mean length and branching of neurons in the ipsilateral somatosensory cortex at day 3 – a defect not observed anymore at the 21 d time point studied for comparison (Hu et al., 2020). Also three repetitive VEGFD applications over 48 h leading to a net dendritic growth of more than a third within 5 days post MCAO (Mauceri et al, 2020) are well in line with the observed net dendritic growth rates in our study.

In this respect, it is also very interesting that post photothromic ischemia *in vivo* imaging showed a massive remodeling of dendrites in the somatosensory cortex at 2 weeks after stroke but no net growth (Brown et al. 2020). This may suggest that the massive induction of net dendritic growth and the branch induction we observed when studying much earlier time points (day 2-4) actually still continue after reaching net recovery at day 4 but that at later stages of cortical remapping are marked by counteraction by dendritic pruning processes operating in parallel.

Such a biphasic repair process would also explain apparently different findings concerning the effects within the dendritic arbor. During the day 2-4 time window of Cobl-dependent acute ischemic stroke-induced repair, all aspects of dendritic arborization were promoted with equal efficiency. The induced repair also did not discriminate between apical and basal dendrites or between the proximal and distal arbor but promoted neuronal morphology in all dendritic areas equally well (this study). In contrast, later dendritic dynamics did not result in any net growth anymore, as dendritic pruning processes operating in parallel lead to higher net growth away from the infarct area and to a net loss of dendrites oriented towards the infarct area (Brown et al., 2020). Together, such a biphasic response would very effectively remap neuronal circuits in the neighborhood of infarct areas.

The observed readdition of on average 5-6 dendritic branching points and of about 400 µm of dendritic arbor per neuron in peri-infarct areas in a time window of repair, which opens at or after day 2 after ischemic stroke, obviously provides a powerful mechanism for not only reconnecting cells that anyway are close enough for synapse formation but also allows for making connections over larger ranges. Consistently, cortical remapping is an important mechanism in recovery after ischemic stroke (Nudo et al., 1996).

Neurons in the penumbra of *Cobl* KO mice subjected to ischemic stroke failed to show any post-ischemic repair. Even 4 and 7 days after MCAO, all quantitative parameters of the dendritic arbor of both layer II/III and layer V neurons in M1 still remained severely affected irrespective of whether the complete dendritic arbor was analyzed or apical and basal dendrites were evaluated separately, whereas WT mice accomplished a full dendritic regrowth in a time frame of about three days following the most severe defects at 24 h post stroke.

What is so special about the actin nucleator Cobl that it is employed in repair processes aiming at regaining brain functions after ischemic stroke in the cortex? Cobl is a powerful actin nucleator that can even work with very low levels of ATP-loaded G-actin, as demonstrated in reconstitutions of actin nucleation with as little as 2 µM actin (Ahuja et al., 2007). Unlike e.g. the Arp2/3 complex, which represents a major actin nucleator for many basic functions of mammalian cells (Pizarro-Cerdá et al., 2016), Cobl is an evolutionary relatively young nucleator with rather specialized functions in a distinct set of specialized cells. These include embryonic hippocampal neurons (Ahuja et al., 2007), early postnatal Purkinje cells in the cerebellum (Haag et al., 2012) and, as demonstrated here in adult mice, also layer II/III and V pyramidal neurons in peri-infarct areas after ischemic stroke. In addition to neurons, they also include early postnatal outer hair cells in the cochlea (Haag et al., 2018) and short-lived enterocytes in the small intestine in mice (Beer et al., 2020) as well as motile cilia-carrying cells in the Kupffer’s vesicle (Ravanelli and Klingensmith, 2011) and sensory stereocilia- and kinocilia-bearing neuromasts of the lateral line organ (Schüler et al., 2013) in fish larvae. In both outer hair cells in the postnatal cochlea and in enterocytes of the small intestine, Cobl seems to be responsible for a specialized set of actin filaments (Haag et al., 2018; Beer et al., 2020). At nascent dendritic branch initiation sites in embryonic neurons, Cobl and F-actin accumulated shortly before and during the branch induction step that breaks the cylindrical symmetry of the respective dendrite segment (Hou et al., 2015). With such specialized cytoskeletal functions, Cobl can thus likely be specifically applied in distinct cellular functions without affecting basic functions of the actin cytoskeleton of a given cell.

A prerequisite for such a molecular scenario of course would be that Cobl’s activity and membrane targeting can be regulated in a distinct manner. Indeed, Cobl functions require arginine methylation of one of its three G-actin binding WH2 domains by the arginine methyltransferase PRMT2 (Hou et al., 2018). Furthermore, Cobl’s N terminal membrane-associating domain (the Cobl Homology domain) interacts with syndapin I in a Ca^2+^/CaM-regulated manner in early neuromorphogenesis (Hou et al., 2015) and links Cobl to its functional partner Cobl-like (Izadi et al., 2021). Strikingly, also Cobl’s actin nucleating, WH2 domain-containing C terminal part was found to be controlled by association of Ca^2+^-activated CaM (Hou et al., 2015).

As unveiled here, Ca^2+^ can also have very detrimental effects on the fate of Cobl and can lead to proteolytic destruction of Cobl. These ying and yang effects of Ca^2+^ on the functions of Cobl are reminiscent of the concept of neurohormesis, in which neurotransmitter stimuli at low and medium intensity are beneficial and lead to functional adaptation, such as synaptic plasticity, whereas high loads of neurotransmitter lead to excitotoxicity (Mattson and Cheng, 2006; Calabrese, 2008). NMDA receptor-mediated Ca^2+^ influx is an important trigger in glutamate-induced neuroexcitotoxicity (Ankarcrona et al., 1995; Glazner et al., 2000; Jaenisch et al., 2016). Consistently, we found that the proteolytic degradation of the actin nucleator Cobl in cortical neurons i) could be induced by incubation with glutamate, ii) was mediated by Ca^2+^ and by calpain and iii) required the opening of NR2B subunit-containing NMDA receptors.

In line with an acute demand of Cobl for dendritic arbor repair after ischemic stroke, Cobl protein levels lost upon the Ca^2+^/calpain-mediated proteolysis were restored to normal levels already 24 h after MCAO. The discovery of increasing *Cobl* mRNA levels during the hours prior to this restoration of Cobl protein levels strongly suggests that this replenishment of Cobl levels is achieved by the observed rise in mRNA level. In line with a decline of Cobl protein levels in M1, also the transient elevation of *Cobl* mRNA levels could be detected in the M1, i.e. in the prenumbra. While nothing is known yet about how the expression of Cobl is controlled, it seems likely that the mRNA increase will be linked to signaling cascades triggered by high glutamate, such as massive increases in cytosolic and thereby also nuclear Ca^2+^ concentrations.

The Ca^2+^/calpain-mediated destructions of the important postsynaptic scaffold protein PSD95, of spectin/fodrin and of the NR2B subunit of NMDA-type glutamate receptors during the first hours after ischemic stroke received great attention, as they can be considered as being linked to the loss of synaptic connections (Simpkins et al., 2003; Pike et al., 2004; Gascón et al., 2008). Surprisingly, none of these components showed any signs of rapid counteraction by increase of mRNA levels. Thus, the transient increase of *Cobl* mRNA coinciding with subsequent restoration of normal Cobl protein levels and initiation of dendritic arbor repair seemed to be a unique response to ischemic conditions. The changes in *Cobl* mRNA levels and the subsequent restoration of Cobl protein levels we observed furthermore are i) consistent with each other and ii) in line with the urgent requirement of Cobl for dendritic arbor repair after ischemic stroke.

Taken together, our study unveiled that ischemic stroke causes damages in dendritic arborization in peri-infarct areas, which then are repaired by processes of dendritic regrowth relying on the actin nucleator Cobl, whose levels upon ischemic stroke first are negatively affected but then are rapidly restored. The excitotoxicity-induced degradation of Cobl was caused by glutamate and NMDA receptor-mediated Ca^2+^ influx and calpain-mediated proteolysis. Subsequent to a rapid restoration of normal Cobl levels, the actin nucleator then powered dendritic repair during a time window opening between day 2 to 4 after ischemic stroke. We show that Cobl-dependent post-stroke dendritic arbor regrowth is a powerful cell biological process that adds to the well-known structural plasticity of post-synapses/dendritic spines.

Importantly, Cobl-dependent post-stroke dendritic arbor regrowth represents a much more long-range mechanism of post-stroke repair inside of peri-infarct areas.

The high conservation of Cobl between mice and man suggests that related Cobl-dependent processes of repair after ischemic stroke may also exist in human patients and would be worthwile to exploit in the identified post-stroke time window.

## Material and Methods

### Experimental Model and Subject Details

#### E. coli strain XL10 Gold

E. coli strain XL10 Gold (Stratagene/Agilent) (genomic information: endA1 glnV44 recA1 thi-1 gyrA96 relA1 lac Hte Δ(mcrA)183 Δ(mcrCB-hsdSMR-mrr)173 tetR F’[proAB lacIqZΔM15 Tn10(TetR Amy CmR)]) was used for amplifications of DNA plasmids. The strain was grown in LB medium (Carl Roth gmbH & Co.KG) or on LB-agar Carl Roth GmbH & Co.KG) with or without the respective antibiotics the strain carries resistances for (10 μg/ml tetracycline, 30 μg/ml chloramphenicol) or obtained resistances for (by transformation with plasmids) (usually either 100 μg/ml ampiciline or 25 μg/ml kanamycine). The strain was usually either grown at RT or at 37°C.

### HEK293 cells

HEK293 cells (Human Embryonic Kidney 293) were cultured in Dulbecco’s modified Eagle’s medium supplemented with 10% fetal bovine serum, 0.1 mM gentamicin, penicillin (100 units/ml) and streptomycin (100 μg/ml) in an cell incubator (5% CO_2_, 37°C, 90% humidity) and split every 3-4 days.

The sex of the cells is not specified. The original publication reports that “primary or early passage secondary human embryonic kidney (HEK) cells prepared by standard techniques” were used (Graham et al., 1977).

### Rat Primary Neurons

Rat cortical neurons (rat strain, Wistar) were prepared as described (Wolf et al., 2019) and cultured until DIV15.

### Mice

All studies were performed with 3-4 months old male WT and *Cobl* KO mice, respectively. *Cobl* KO mice were described previously (Haag et al., 2018). They were generated via excision of exon 11 of the *Cobl* gene using the Cre/loxP system and subjected to speed congenics (Haag et al., 2018). For the current study, *Cobl* KO mice were backcrossed to C57BL/6J until C57BL/6J::129/SvJ (97.375::2.625) was reached (infarct volumetry) (B6.129/Sv-Cobl^tm1BQ^). Neuromorphometric analyses were mostly done with even further backcrossed *Cobl* KO animals (upto 99.918% C57BL/6J). Despite the high C57BL/6J genetic backgrounds, comparisons with WT animals were done with WT littermates of heterozygous breedings of *Cobl* KO mice in both infarct volumety and morphometric analyses of stroke damage and repair processes.

Additionally, tissue material for qPCRs and Western blot analyses as well as for brain sections for anti-MAP2 and Golgi-Cox stainings were also taken from WT C57BL/6J mice when MCAO conditions were examined and compared in different WT conditions.

## Methods

### Breeding, housing and handling of mice

Mice were kept in a specific pathogen-free environment at room temperature (22°C) with the humidity of 68% under 14 h light/10 h dark conditions with *ad libitum* access to food and water. Mice were regularly tested for parasites and other pathogens using sentinel mice held under identical conditions.

All animal care and experimental procedures were performed in accordance with EU animal welfare protection laws and regulations and were approved by a licensing committee from the local government (Landesamt für Verbraucherschutz, Bad Langensalza, Thuringia, Germany). The generation of *Cobl* KO mice was approved by the permission number 02-011/10. Breeding and housing was approved by the permission number UKJ-17-021. MCAO experiments with WT mice were approved by permission numbers 02-024-15 and 02-057-14, respectively. The MCAO experiments conducted with WT and *Cobl* KO littermates from heterozygous breedings of *Cobl* KO mice were approved by the permission number UKJ-18-030 (Landesamt für Verbraucherschutz, Bad Langensalza, Thuringia, Germany).

### MCAO

MCAO can be experimentally achieved in mice and rats by temporarily occluding the common carotic artery (CCA), introducing a suture directly into the internal carotic artery (ICA) and advancing it until it interrupts the blood supply to the MCA (Popp et al., 2009; Heiss, 2000). MCAO therefore is one of the most frequently used experimental ischemic stroke model systems. 30 min of ischemia causes neuronal cell death limited to the striatum. MCAO was induced as previously described (Sieber et al., 2010). In brief, mice were treated with Melosus and anesthetized with 2.5% (v/v) isoflurane in N_2_O:O_2_ (3:1). Through a middle neck incision, the right CCA, the external carotid artery (ECA) and the ICA were carefully dissected from surrounding nerves and fascia. A 7-0 nylon monofilament (70SPRe, Doccol Corp.) coated with silicone rubber on the tip (0.20 ± 0.01 mm diameter) was introduced into the ICA through an incision of the right CCA up to the circle of Willis. In this position the suture occluded MCA. During the MCA occlusion body temperature was maintained at physiological level using a heating pad. After 30 min of occlusion the suture was withdrawn to restore the blood flow and allow for reperfusion. Sham animals underwent anesthesia and the same surgical procedures except the occlusion of MCA.

### Golgi-Cox staining, cryosectioning and infarct validation

Golgi-Cox staining (Golgi-Cox impregnation) for studying the morphology of individual neurons in the brain (Das et al., 2013) was done with brain material from adult (3-4 months old), male mice that were subjected to either sham or 30 min MCAO and sacrificed after different reperfusion times. The brains were quickly removed and separated into two parts using a Precision Brain Slicer (Braintree Scientific Inc.) at bregma +0.8 mm using the Allen Brain Atlas as reference. The anterior part was used for the 25 infarct validation by anti-MAP2 staining (see below), and the posterior part was used for Golgi-Cox staining using FD Rapid GolgiStain kits (FD NeuroTechnologies, Inc), essentially following a protocol described previously (Schneider et al., 2014; Koch et al., 2020). In brief, the posterior brain part was immersed in solution A+B of the FD Rapid GolgiStain kit and kept in the dark at RT for 21 days. Brains were then incubated in solution C of the kit for another 5 days (RT; dark). The brains were then dipped very slowly into dry ice-precooled isopentane and stored at -80°C.

Cryosectioning was performed at -28°C using a Leica CM 3050 S (Leica Biosystems Nussloch GmbH). Coronal sections of 100 µm thickness were cut and transferred onto small drops of solution C of the FD Rapid GolgiStain kit on gelatine-coated slides. After drying overnight (RT; dark), the sections were rinsed twice with water and stained for 1 min in the staining solution of the kit. After rinsing twice with water, sections were dehydrated in an ascending ethanol series (50%, 75%, 95%, 4times 99% (v/v)) and cleared three times in xylene. Afterwards, the Golgi-Cox-stained tissue sections were mounted onto coverslips with Roti-Histokitt (Carl Roth GmbH & Co. KG) and stored in the dark.

The anterior part of the brains cut at bregma +0.8 mm was used for anti-MAP2 immunostainings. For this purpose, free-floating sections were pretreated with 0.24% (v/v) H_2_O_2_ in Tris-buffered saline (83.8 mM Tris-HCl; 16 mM Tris-Ultra (Base) pH 7.4, 154 mM NaCl, 0.2% (v/v) Triton X-100) (TBS-T) for 30 min and blocked with TBS-T containing 3% (v/v) normal donkey serum (NDS) (GeneTex Inc.), 2% (w/v) bovine serum albumin (Sigma Aldrich Co. LLC), 3% (w/v) milk powder and Fab fragments of donkey anti-mouse IgG (1:200) (Jackson Immunoresearch).

Sections were incubated at 4°C overnight with monoclonal antibody against mouse MAP2 (Sigma-Aldrich) (1:1000 in TBS-T with 3% (v/v) NDS) and then with a biotinylated secondary antibody (Jackson ImmunoResearch) (1:500 in TBS-T with 3% (v/v) NDS) after 3 times washing with TBS-T and 1 time with 3% (v/v) NDS. In the end, the sections were processed with Vectastain Elite ABC HRP kits (Vector Laboratories) for 1 h and immunosignals were visualized using 3,3′-diaminobenzidine reactions. Brains of MCAO-treated mice displaying an anti-MAP2-negative infarct area in the ipsilateral hemisphere were then subjected to the neuromorphological analysis and compared to brains of sham-treated animals processed the same way. Blinding was done by using multiple-digit animal numbers not indicative of genotype, MCAO or sham treatment or the different reperfusion times and by using Antrenamer2 blinding software.

### Quantitative analyses of neuronal morphology in the M1

Image acquisitions for neuronal morphology analyses were done from 3-5 independent coronal sections containing the M1 area per mouse (Bregma +0.8 to 0 mm). Layer II/III and layer V pyramidal neurons in M1 were identified by their distance from pia and their distinct morphologies. Z-stacks of Golgi-Cox-stained neurons were recorded at 20x magnification (1 µm intervals), using a Zeiss AxioObserver equipped with a Zeiss Plan-Apochromat 20x/0.5 objective and an AxioCam MRm CCD camera (Zeiss). Digital images were acquired by AxioVision software (Zeiss).

Overview images of mouse brain sections were generated by stitching images recorded with a 5x/0.16 objective together using the Tiles mode of the microscope and the automatic stitching of the Zen Software (Zeiss) with an overlap of 10%.

Three-dimensional reconstructions of neuronal morphologies were performed with Imaris 8.0 software (Bitplane) using the filament tracer set with shortest distance algorithm (diameter: 2-16 µm). The images were baseline- and background-subtracted as described before (Koch et al., 2020). As in previous software-based quantitiative determinations of dendritic arbor complexity evaluations (Izadi et al., 2018) the following parameters were analyzed: the number of Sholl intersections, the number of dendritic branching points, the numbers of dendritic terminal points and the total dendritic tree length.

The basal dendrites and the arbor originating from the apical dendrite were analyzed accordingly by dissecting the Imaris tracings of evaluated layer II/III WT and *Cobl* KO neurons.

The analyses were conducted in a blinded manner. Quantitative parameters were analyzed for data distribution and statistical significances with Prism 6 software (SCR_002798; GraphPad).

### Cobl gain- and loss-of-function studies in primary rat cortical neurons during early development

Rat cortical neurons were prepared and transfected with plasmids (GFP and GFP-Cobl for gain-of-function experiments; scrambled (scr.) RNAi/GFP (control) and Cobl RNAi/GFP (Ahuja et al., 2007) as well as Cobl RNAi/GFP-Cobl* (rescue; Haag et al., 2012) for loss-of-function experiments) at DIV4, as described (Wolf et al., 2019). Neurons were fixed 40 hours after transfection by 4% (w/v) paraformaldehyde (PFA) for 7 min, then permeablilized and blocked by blocking solution consisting of 10% (w/v) horse serum, 5% (w/v) bovine serum albumin and 0.2% (v/v) Triton X-100 in PBS for 1 hour at room temperature. After the immunostaining with anti-MAP2 antibodies (secondary antibodies, Alexa Fluor 568-labelled donkey anti-mouse (Molecular Probes (A10037)), the transfected neurons from two independent coverslips per condition per assay were imaged using a Zeiss AxioObserver Z1 microscope/ApoTome.

Morphometric measurements were based on the MAP2 signals using Imaris 8.4.0 software. The detailed settings were: thinnest diameter and gap length: 2 µm, minimum segment length: 10 µm. The number of dendritic branching points, terminal points, total dendritic tree length and Sholl intersections were analyzed as described above for Golgi-stained neurons in M1.

### Determination of infarct volumes

The volumes of infarct areas after 30 min MCAO were determined after 7 days of reperfusion according to procedures described previously (Stahr et al., 2012). In brief, *Cobl* KO and WT mice were anesthetized by isoflurane and transcardial perfusion was performed with 4% (w/v) PFA in 0.2 M sodium phosphate buffer pH 7.4. Brains were then removed and post-fixed in 4% (w/v) PFA for 5 h, cryoprotected in 0.1 M sodium phosphate buffer (pH 7.4) containing 10% (w/v) sucrose overnight and transferred to 30% (w/v) sucrose in 0.1 M sodium phosphate buffer (pH 7.4) until saturation. After freezing, the brains were stored at -80°C.

Coronal sections (40 µm) were sliced with a cryomicrotome (Thermo Fisher Scientific Inc.). The sections were maintained in anti-freeze buffer (0.05 M sodium phosphate buffer (pH 7.4) containing 3 mM NaN_3_, 832 mM glucose and 30% (v/v) ethylene glycol) at -20°C until use. Free-floating sections were subjected to anti-MAP2 immunostainings as described above.

The anti-MAP2 stained brain sections were imaged on a light table using a digital CCD camera (Hamamatsu Photonics) and the sizes of the infarct area as well as of each hemisphere and of the total brain section were measured by Scion Image software (Scion Corporation). The area of the infarct was traced and quantified on every twelfth section. Infarct volumes were calculated as percentage of the total brain.

### Brain tissue preparations for subsequent qPCR and Western blot analyses

Mice subjected to 30 min MCAO were decapitated under deep anesthesia at 2 h, 3h, 6 h, 12 h, 24 h and 48 h, respectively, reperfusion time. Brains were removed and tissue samples for expression analyses were prepared as described previously (Sieber et al., 2010). In brief, brains were cut into three coronal segments using a Precision Brain Slicer (Braintree Scientific Inc.): from +2.8 to +0.8 mm to bregma (rostral segment), from +0.8 to -1.2 mm to bregma (middle segment), and from -1.2 to -3.2 mm to bregma (caudal segment). The ipsilateral part (ischemic hemisphere) and the contralateral part (non-ischemic hemisphere) of the middle brain segment were separated and quick frozen in liquid nitrogen and stored at -80°C for later processing.

In further animal experiments (after 6 h, 12 h and 24 h reperfusion time), the middle brain segment was dissected further and the primary motor cortex M1 was isolated and snap-frozen for subsequent analyses of exclusively penumbral ipsilateral and contralateral brain areas.

### qPCR analyses

Slices from rostral and caudal segments that were adjacent to the middle segment were subjected to anti-MAP2 staining for infarct validation to select mice with similar lesion size for mRNA quantifications, as described before (Jaenisch et al., 2016).

In brief, the ipsilateral and contralateral parts of the middle brain segment and M1 tissue samples of mice with comparable MCAO-induced lesions prepared as described above were homogenized in QIAzol Lysis Reagent (Qiagen GmbH) followed by the addition of chloroform (Sigma Aldrich Co. LLC) to separate the homogenate into a clear upper aqueous layer (containing RNA), an interphase, and a red lower organic layer (containing the DNA and proteins). RNA was precipitated from the aqueous layer with isopropanol. The precipitated RNA was washed with 75% (v/v) ethanol and then resuspended in pure H_2_O (Gibco). The quality and quantity of the RNA were measured by Nanodrop 2000 (Thermo Fisher Scientific Inc.).

After adjustment of RNA concentration, equal amounts of total RNA (0.5 µg) were transcribed to cDNA with RevertAid First Strand cDNA synthesis kit (Thermo Fisher Scientific Inc.). In detail, 0.6 µl oligo(dT)18 primer (stock, 100 µM) and 0.5 µl random hexamer primer (stock, 100 µM) were mixed and added to 5 µl RNA (100 ng/µl), and then incubated in a thermal cycler for 5 min at 65°C. Meanwhile, 2 µl 5x reaction buffer, 1 µl dNTP (10 mM), 0.5 µl RiboLock RNase inhibtor (20 U/µl) and 0.5 µl Revert Aid reverse transcriptase (200 U/µl) were mixed and then added to the pre-incubation-mix. Transcription was done by incubations at 25°C (5 min), at 42°C (60 min) and finally at 70°C (5 min). The obtained cDNA was then diluted to the final concentration of 5 ng/µl and used for qPCR analyses.

qPCR was performed in a volume of 20 µl containing Brilliant III SYBR® Green qPCR Master Mix (Agilent), cDNA (equivalent to 25 ng reversely transcribed RNA) and specific primers each at a final concentration of 500 nM. Primers used are listed in Table 1.

**Table 1.**
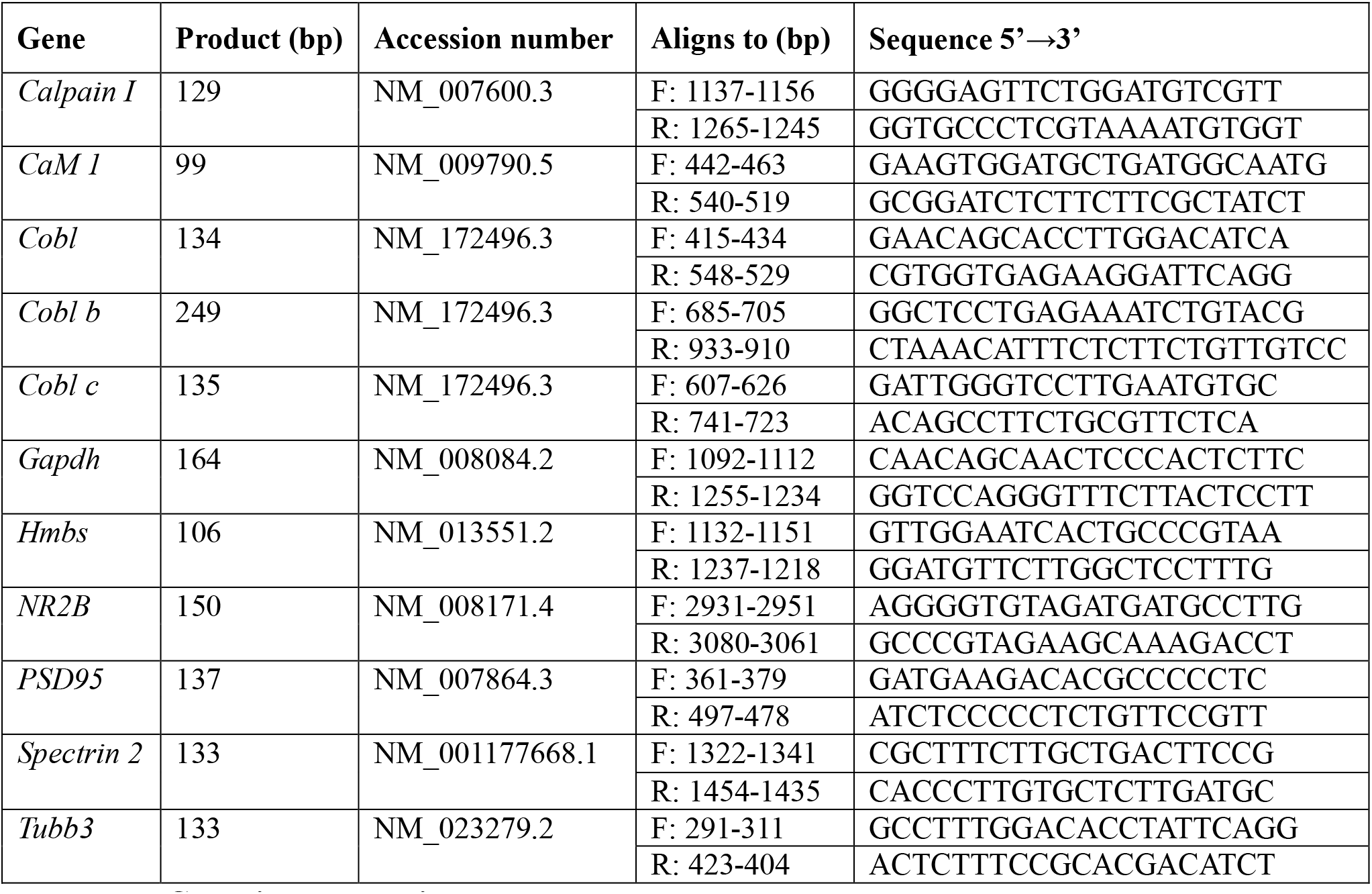
qPCR primers used in the study.

Amplification was performed using a Rotor-Gene 6000 (Qiagen GmbH) (cycle conditions: 3 min polymerase activation, 40 amplification cycles of 95°C for 10 s and 60°C for 15 s each).

Gene expression data were normalized to *glyceraldehyde 3-phosphate dehydrogenase* (*Gapdh*), *hydroxymethylbilane synthase* (*Hmbs*) and *tubulin beta 3 class III* (*Tubb3*) data, respectively (*Gapdh*, *Hmbs* and *Tubb3* primers, see Table 1). Relative expression levels were calculated using the Pfaffl equation (Pfaffl, 2001), 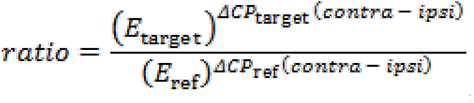, with *E*_target_ being the real-time PCR efficiency of target gene amplification, *E*_ref_ being the real-time PCR efficiency of a reference gene transcript, ΔCP_target_ being the CP deviation of contra minus ipsi data of the target gene transcript and ΔCP_ref_ being the CP deviation of contra minus ipsi data of the reference gene.

### Protein isolation from mouse brain tissue for quantitative immunoblotting analyses

The ipsilateral part and the contralateral part, respectively of the middle brain segment (+0.8 and -1.2 mm to bregma) as well as ipsilateral and contralateral M1 tissue samples were prepared as described above, homogenized and immunoblotted.

In order to improve comparability, samples of the same condition were analyzed on the same blotting membrane. Linearity of the signals was ensured by using fluorescence-based immunodetections analyzed by a LI-COR Odyssey System (LI-COR Bioscience).

Primary antibodies used included polyclonal guinea-pig anti-Cobl antibodies (DBY; WB, 1:500) (Haag et al., 2012), monoclonal mouse anti-actin antibodies (AC-15; Sigma; #A5441; WB, 1:5000), polyclonal rabbit anti-ß3-tubulin antibodies (Synaptic Systems; #302 302; WB, 1:2000) and monoclonal mouse anti-fodrin antibodies (clone AA6; Enzo Life Sciences; #BML-FG6090; WB, 1:4000).

Secondary antibodies used were DyLight800-conjugated goat anti-rabbit and anti-mouse antibodies (Thermo Fisher Scientific; #SA5-35571, #SA5-35521) as well as donkey anti-guinea pig antibodies labelled with IRDye680 and IRDye800 (LI-COR Bioscience; #926-68077, #926-32411). For comparison of contra vs. ipsi, the contra signal was set to 100%. The ipsi signal was expressed as percent of contra signal (of the same brain sample).

For examinations of Cobl’s Ca^2+^- and calpain-dependent proteolysis, mouse brain homogenates were generated in lysis buffer (10 mM HEPES pH 7.4, 1% (v/v) Triton X-100, 0.1 mM MgCl_2_) containing 10 mM NaCl and 1x Complete protease inhibitor without EDTA (Roche) and were then either treated with EDTA (1 mM) or incubated with 100 µM Ca^2+^ with and without calpain inhibitor I (CP-1; 15 min, 4°C), respectively, and subsequently subjected to anti-Cobl and anti-actin immunoblotting analyses.

### Treatments of primary rat cortical neurons and protein isolation from such cultures for quantitative immunoblotting analyses

Rat cortical neurons were prepared as described (Wolf et al., 2019) and cultured until DIV15.

For glutamate stimulations, the primary neuronal cell cultures at DIV15 were washed with Hanks’ Balanced Salt Solution (HBSS), and the initial medium was replaced by medium containing 40 µM glutamate. Examined were the excitotoxicity effects induced by 0, 5, 10, 30 and 60 min of incubation with glutamate (compare also Ankarcrona et al. (1995) and Xu et al. (2009)).

In some experiments, inhibitors and antagonists (Calpeptin, 20 µM; CP-1, 20 µM; Chloroquine, 500 µM; Lactacystin, 5 µM; MK801, 50 µM; Ifenprodil, 10 µM; CNQX, 40 µM; AP-5, 100 µM final concentration) were added to the medium (30 min preincubation; according to Xu et al. (2009)). After the preincubations, the cells were then treated with 40 µM glutamate for 30 min.

After stimulation, cells were scraped off in lysis buffer containing 10 mM NaCl, 1x Complete protease inhibitor without EDTA (Roche) and 1 mM EDTA. The lysates were sonicated (2 pulses), the homogenates were centrifuged for 10 min (4°C, 1000 x g) and the protein content was precipitated (20% aceton overnight) at -20°C. The protein pellet was collected by centrifugation (20000 x g, 4°C, 10 min), dried, resuspended in 1x SDS sample buffer and incubated for 10 min at 75°C. The samples were then subjected to Western blotting using antibodies described above as well as monoclonal mouse antibodies against PSD95 (6G6-1C9; Abcam; #ab2723; WB, 1:2000), NR2B (13/NMDAR2B; BD Bioscience; #610416; WB, 1:500), and Arp3 (Abcam; #ab49671; WB, 1:1000).

The immunosignals were analyzed quantitatively with a LI-COR Odyssey System.

### In vitro reconstitution of Cobl proteolysis by calpain I

HEK293 cells were transfected with plasmids encoding for GFP-Cobl^1-713^ and GFP-Cobl^712-1337^, respectively, which were generated by subcloning from GFP-Cobl full length (Ahuja et al., 2007) using an internal Hind III site, and with GFP-Cobl^1-408^, GFP-Cobl^406-866^, GFP-Cobl^750-1005^, GFP-Cobl^1001-1337^ (Ahuja et al., 2007; Schwintzer et al., 2011; Hou et al., 2015). One day after transfection, HEK293 cells were washed with PBS, harvested and lyzed by incubation in lysis buffer containing 75 mM NaCl, 1x Complete protease inhibitor without EDTA and 1 mM EGTA for 30 min at 4°C. Cell lysates were obtained as supernatants from centrifugations at 20000 x g (20 min at 4°C).

Anti-GFP antibodies (ab290; Abcam) immobilized to protein A-agarose (Santa Cruz Biotechnology; #sc-2001) were used to immunocrecipitate the GFP-fusion proteins. After incubation with the HEK293 cell lysates for 3 h at 4°C, anti-GFP antibody-associated proteins were isolated by centrifugation at 11000 x g for 1 min and washed 2 times with lysis buffer containing 75 mM NaCl. One sample was incubated in cleavage buffer (PBS pH 7.4, 1 mM L-cysteine, 2 mM CaCl_2_), whereas the others were washed once with cleavage buffer and then incubated at 25°C with calpain I in 50 μl cleavage buffer for 10 min (final concentrations, 0, 0.001, 0.01 and 0.1 U/µl). The cleavage reactions were stopped by adding 4×SDS sample buffer and by denaturing at 95°C for 5 min.

The proteolytic products were immunoblotted with monoclonal mouse anti-GFP antibodies (JL8; Clontech; RRID: AB_10013427) and analyzed by a LI-COR Odyssey System.

### Quantification and statistical analysis

All quantitative data shown represent mean±SEM and are displayed with overlayed dot plots wherever useful. Tests for normal data distribution and statistical significance analyses were done using Prism 6 software (GraphPad; SCR_002798).

Statistical significances of neuronal morphological analyses in Golgi-Cox-stained M1 tissue samples as well as Cobl-loss-of function and Cobl gain of function data in cultures of cortical neurons were tested using Mann-Whitney, one-way ANOVA with Tukey’s post test and two-way ANOVA with Sidak’s post-test, and Student’s t-test, respectively. Statistical analyses of Sholl intersections were performed using two-way ANOVA with Sidak’s post-test.

Wilcoxon signed rank test was used for changes detected in Cobl protein levels in post-ischemic brains. One-way ANOVA with Dunn’s post-test was applied to biochemical analyses of Cobl degradation in examinations of glutamate-induced excitotoxicity and the molecular mechanisms involved.

Relative mRNA expression levels in post-ischemic brains and in sham animals were tested for one-way ANOVA with Sidak’s post test.

Infarct volumes were statistically analyzed by unpaired, two-tailed Student’s t test.

## Contact for Reagent and Resource Sharing

Further information and requests for resources and reagents should be directed to the Lead Contact, Michael M. Kessels.

## Acknowledgements

We thank I. Ingrisch and M. Öhler for excellent technical support and L. Schwintzer, M. Izadi as well as N. Haag for their generous support with plasmids, practical help and mice, respectively.

This work was supported by grants from the DFG (*Deutsche Forschungsgemeinschaft*) to M.M.K. (KE685/3-2) and to B.Q. (RTG1715) as well as by the IZKF (*Interdisziplinäres Zentrum für klinische Forschung des Universitätsklinikums Jena*) to B.Q. and C.F. (RTG1715 SP18).

## Author contributions

Y.J and D.K. generated and evaluated the data displayed. Y.J. co-wrote the manuscript. Y.J., D.K., B.Q. and M.M.K. designed and/or made figures. M.G. helped with MCAOs and sample preparations and furthermore established methods. J.G.D. carried responsibilities for *Cobl* KO mice backcrossings, breedings, number coding and administrative mouse work. O.W.W. provided scientific advice and resources. M.M.K., C.F. and B.Q. supervised the project, evaluated data and beared responsibilities for mice breedings and animal experiments. M.M.K. and B.Q. wrote the paper.

## Data availability statement

This study includes no data deposited in external repositories.

Accession codes and unique identifiers are given wherever useful.

The following figures have associated raw data: Fig. 1-6 and Fig. S1-5 have associated numerical data; Fig. 1 and Fig. 2 furthermore have uncropped Western blot images as associated raw data (please see Appendix).

## Supplementary information

### Supplementary figures and legends

**Fig. S1 (related to Fig. 2).**
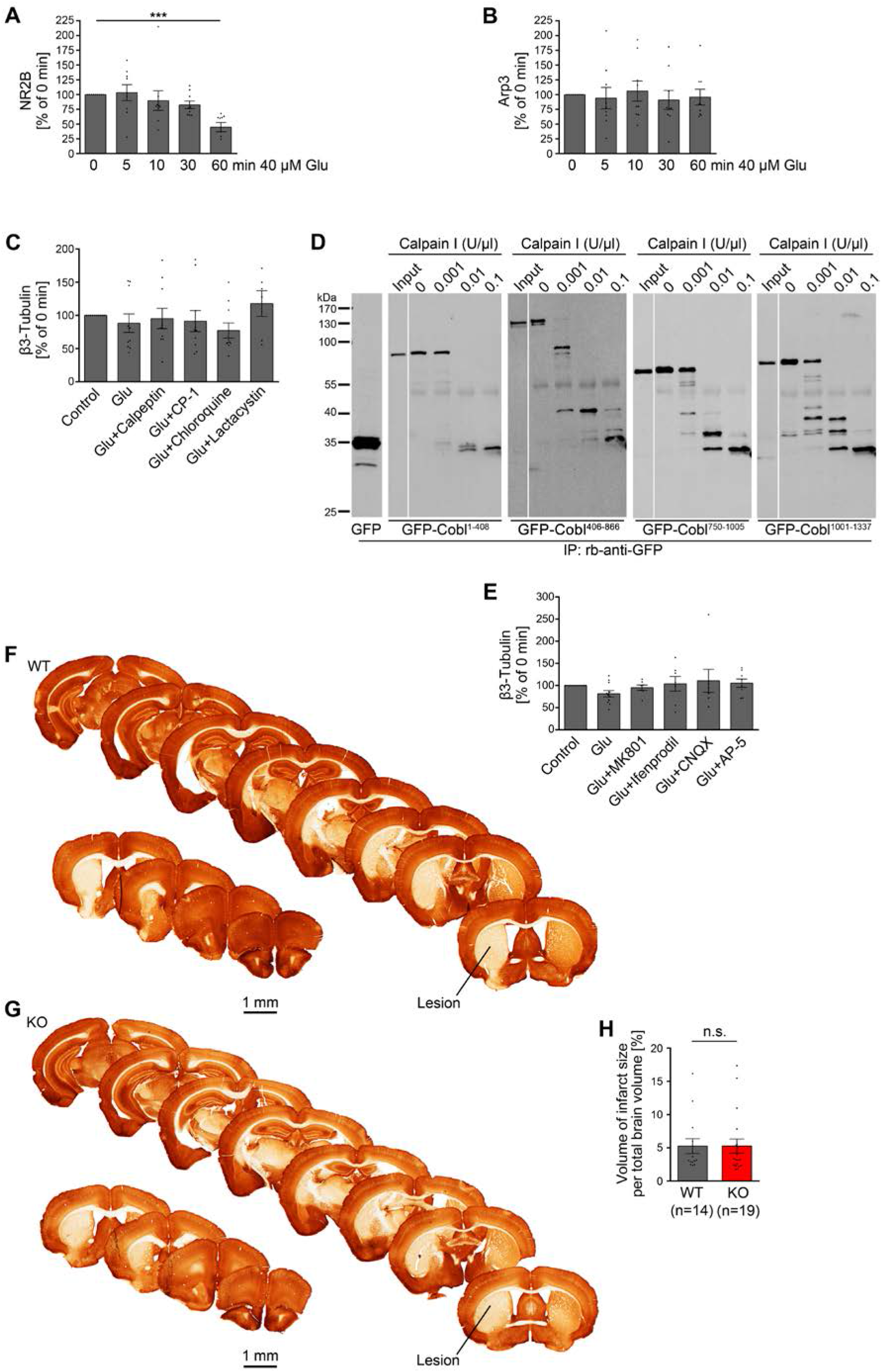
The actin nucleator Cobl is degraded by the Ca^2+^-controlled protease calpain during NMDAR-mediated excitotoxicity but *Cobl* KO does not affect the size of the final lesion caused by ischemic stroke induced by MCAO. **A,B,** Quantitative immunoblotting analyses of NR2B (**A**) and Arp3 (**B**) in cortical neuronal cultures subjected to different durations of incubation with 40 µM glutamate (Glu). n=9 independent assays and biological samples. **C,** Proteolytic pathways underlying the Cobl decline upon prolonged stimulation with glutamate, as shown by the use of inhibitors against calpain (Calpeptin, CP-1), against lysosomal degradation (Chloroquine) and aginst proteasomal proteolysis (Lactacystin), respectively, in quantitative immunoblotting analyses. n_Control_=10, n_Glu_=10, n_Glu+Calpeptin_=10, n_Glu+CP-1_=10, n_Glu+Chloroquine_=10, n_Glu+Lactacystin_=6 independent biological samples. **D,** Anti-GFP immunoblotting analyses of GFP-Cobl^1-408^, GFP-Cobl^406-866^, GFP-Cobl^750-1005^ and GFP-Cobl^1001-1337^ expressed in HEK293 cells, immunoisolated with anti-GFP antibodies and incubated without and with calpain I (10 min, 25°C). Input shows the GFP fusion protein prior to the incubation with calpain concentrations ranging from 0 to 0.1 U/µl. GFP is shown for size comparison. White lines indicate lanes omitted from the blots. Size standards apply to all blots shown. **E,** Lack of effects of inhibitors against open NMDARs (Glu+MK801), against the NR2B subunits of NMDA receptors (Glu+Ifenprodil), against AMPA and kainate receptors (Glu+CNQX), and against NMDARs (Glu+AP-5), respectively, when compared to control and glutamate-induced (30 min, 40 µM) excitotoxicity in quantitative anti-tubulin immunoblotting analyses of lysates of neuronal cultures. n_control_ = 11, n_Glu_ = 11, n_Glu+MK-801_ = 7, n_Glu+Ifenprodil_ = 7, n_Glu+CNQX_ = 7, n_Glu+AP-5_ = 8 independent biological samples. Data represent mean±SEM. Statistical significance calculations, one-way ANOVA with Dunn’s post-test (**A-C,E**). \*\*\**P*<0.001. **F,G,** Representative examples of serial sections of brains of WT (**F**) and *Cobl* KO (**G**) mice at 7 days of reperfusion, respectively, as they were used for determinations of MCAO-induced lesion volumes in WT and *Cobl* KO mice in relation to the total brain size. The sections were immunostained with anti-MAP2 antibodies to visualize the lesions. Note the large MAP2-negative lesions (examplarily marked in one section) visible in both WT and *Cobl* KO brains. Bars, 1 mm. **H,** Quantitative determination of lesion volumes caused by 30 min MCAO in relation to the total brain volume in percent. Note that there was no significant difference in the volume of MCAO-induced lesions when *Cobl* KO brains were compared to WT brains. Data represent means±SEM presented as bar plots overlayed with all individual data points. n_WT_=14; n_KO_=19 mice. Statistical significances were calculated using Student’s t-test (n.s.).

**Fig. S2 (related to Fig. 3).**
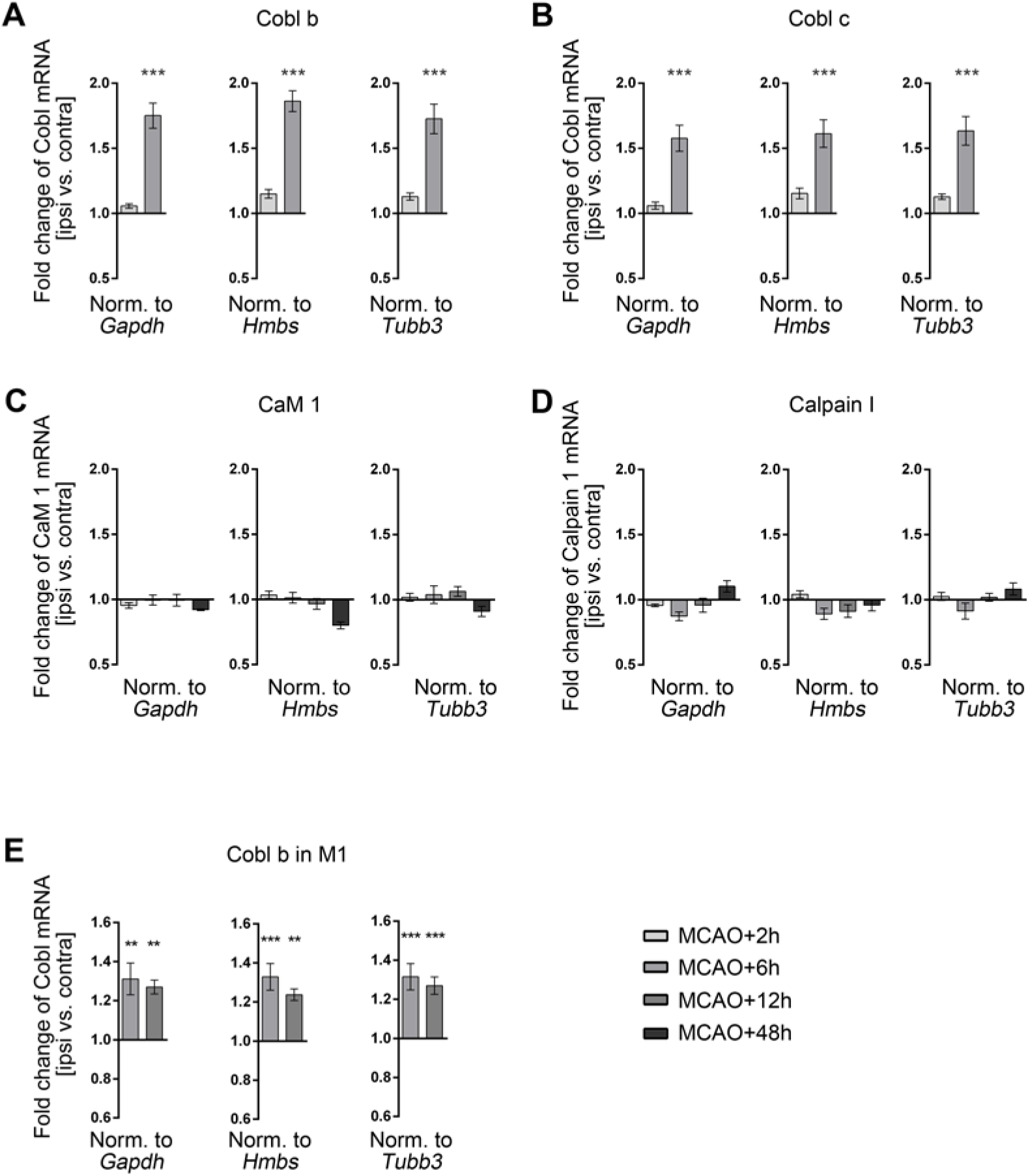
qPCR analyses demonstrate a transient excess of ipsilateral *Cobl* mRNA expression 6 h after MCAO. **A,B,** Fold change of *Cobl* mRNA at 2 h and 6 h reperfusion after MCAO, as determined by qPCR with Cobl b (**A**) and Cobl c (**B**) primers respectively. qPCR data represent the differences between ipsi and contra of *Cobl* (primer sets b and c, respectively) normalized to *Gapdh* (left panel), *Hmbs* (middle panel) and *Tubb3* (right panel), respectively. n _MCAO+2h_=8; n _MCAO+6h_=8 brain samples and mice. **C,D,** Fold change of *CaM 1* (**C**) and *Calpain I* (**D**) mRNA levels at 2 h, 6 h, 12 h and 48 h reperfusion time after 30 min MCAO. Data represent ratios of the differences between ipsi and contra of *CaM 1* and *Calpain I* levels, respectively, normalized to *Gapdh*, *Hmbs* and *Tubb3*, respectively. **E,** Fold changes of *Cobl* mRNA levels in M1 tissue samples (ipsi vs. contra) 6 h and 12 h after MCAO. The data is again normalized against three different genes (*Gapdh*, *Hmbs* and *Tubb3*). **A-D,** n _MCAO+2h_=8; n _MCAO+6h_=8; n _MCAO+12h_=9; n _MCAO+48h_=6 brain samples and mice. **E,** n_MCAO+6h_=7; n_MCAO+12h_=8 M1 samples and mice. Data, mean±SEM. Statistical significances (ipsi vs. contra) were calculated using one-way ANOVA with Sidak’s post-test (**A-E**), respectively. *\*\*P*< 0.001; *\*\*\*P*< 0.001.

**Fig. S3 (related to Fig. 4).**
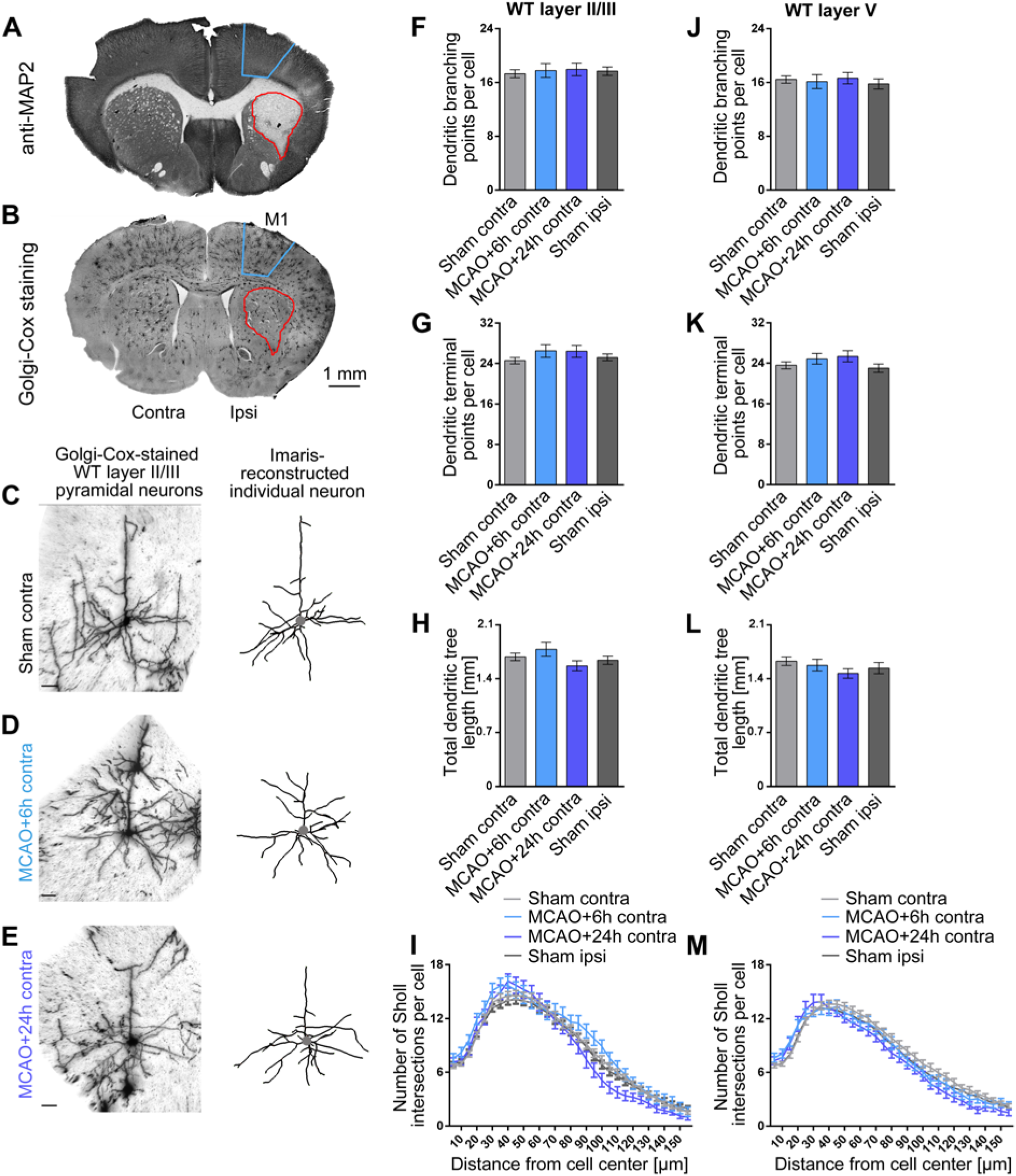
No change in dendritic arbor complexity subsequent to ischemic stroke in layer II/III and layer V neurons in the contralateral motor cortex. **A,B,** Representative micrographs of adjacent serial coronal sections of brains at ∼bregma +0.8 mm from mice (age, 3-4 months) that were subjected to 30 min MCAO. **A**, Anti-MAP2 immunostaining. **B**, Golgi-Cox staining. The lesion caused by the infarct can be determined based on the lack of anti-MAP2 detection (outlined in red). Blue framing indicates the adjacent region M1 used for morphological analysis in corresponding Golgi-Cox-stained sections. **C-E,** Representative images of Golgi-Cox-stained (left panels) and Imaris-reconstructed (right panels) layer II/III pyramidal neurons of M1 from the contralateral side of sham-treated mice (**C**) and of mice subjected to 30 min MCAO analyzed after 6 h (**D**) and 24 h (**E**) reperfusion times. The position of the cell bodies are marked by a gray dot. Scale bars, 30 μm. **F-M,** Quantitative determinations of dendritic branching points (**F,J**), dendritic terminal points (**G,K**), total dendritic tree length (**H,L**) and Sholl intersections (**I,M**) of dendritic trees in layer II/III (**F-I**) and layer V of M1 (**J-M**). Note that all parameters of dendritic complexity at the contralateral side remained unchanged in comparison to sham-treated mice at both 6 h and 24 h after MCAO. Layer II/III: n_Sham contra_=57; n_MCAO+6h contra_=18; n_MCAO+24h contra_=17; n_Sham ipsi_=60 neurons. Layer V: n_Sham contra_=60; n_MCAO+6h contra_=22; n_MCAO+24h contra_=25; n_Sham ipsi_=47 neurons from 3 mice each for the two MCAO conditions (for ipsi data from the same mice see Fig. 4) and from 6 mice for sham controls (contralateral and ipsilateral; sham ipsi data as in Fig. 4 for comparison). Quantitative data represent mean±SEM. Statistical significance calculations, one-way ANOVA with Tukey’s post-test (**F-H,J-L**) and two-way ANOVA with Sidak’s post-test for Sholl analysis (**I,M**), respectively (all n.s.).

**Fig. S4 (related to Fig. 5).**
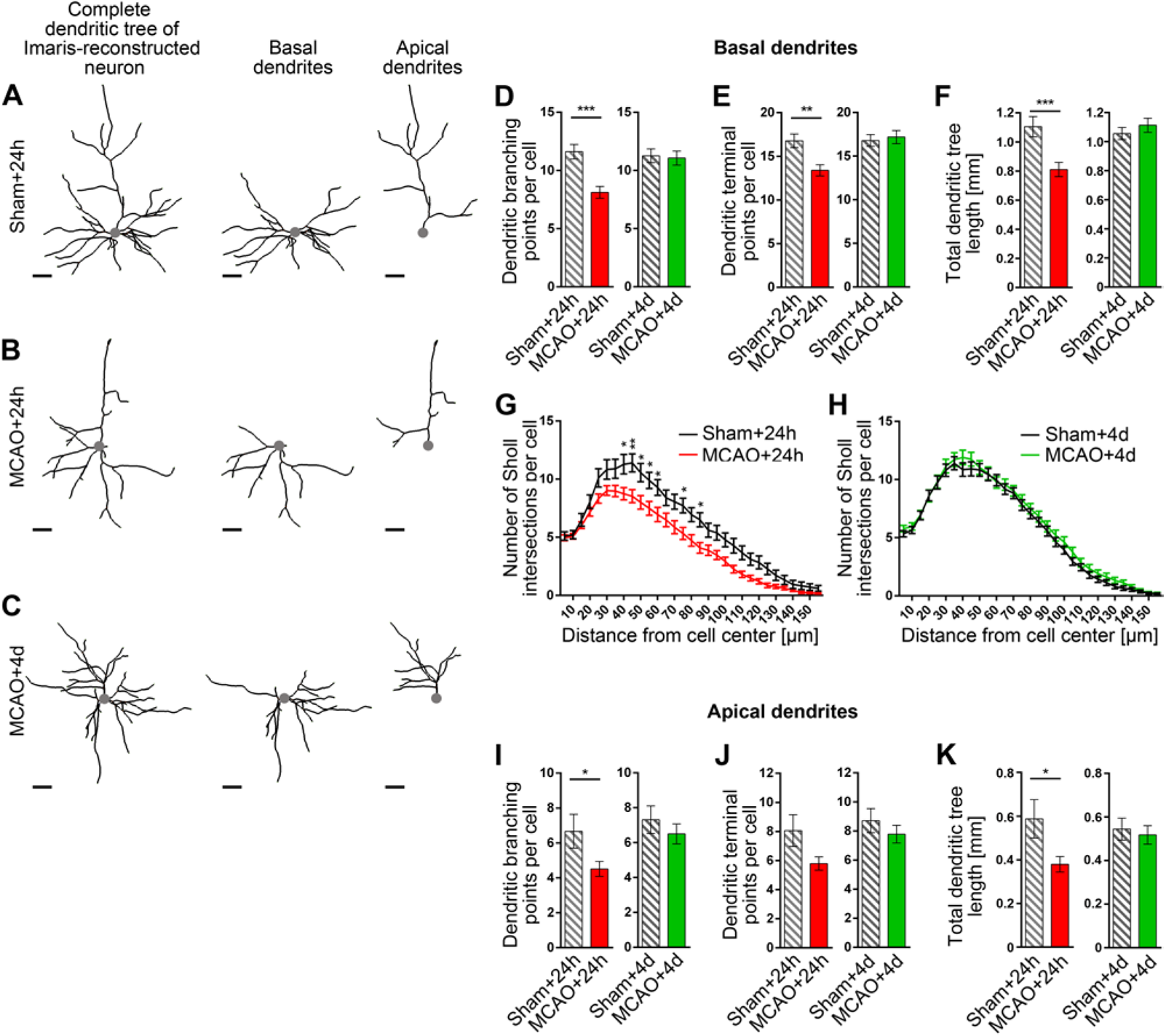
MCAO-induced defects in dendritic arborization manifest in both apical and basal dendrites of layer II/III pyramidal neurons of M1. **A-C,** Individual Imaris-reconstructed cells from Fig. 5 A-C (right panels) showing ipsilateral layer II/III neurons from M1 of WT mice subjected to sham treatment (**A**) and 30 min MCAO with 24 h and 4 d reperfusion time (**B,C**), respectively, for differential analyses of basal (**D-H**) and apical dendritic parameters (**I-K**). The apical and basal dendritic parts are depicted in **A-C**. The position of the cell bodies are marked by a gray dot. Scale bars, 30 μm. **D-K,** Quantitative determinations of dendritic branching points, terminal points, total dendritic tree length and Sholl intersections of basal dendrites (**D-H**) and related analyses for apical dendrites (**I-K**). Note that MCAO-induced defects occurr in both apical and basal dendrites 24 after MCAO and also show full recovery in both apical and basal dendritic arbors after 4 d reperfusion. n_Sham+24h_=18; n_MCAO+24h_ =43; n_Sham+4d_=35; n_MCAO+4d_=32 neurons from 6 mice for the MCAO+24h group and 3 mice for the MCAO+4d group and from 3 mice for each sham control group (ipsi). Quantitative data represent mean±SEM. Statistical significance calculations, Mann-Whitney (**D-F,I-K**) and two-way ANOVA with Sidak’s post-test for Sholl analysis (**E,F**), respectively. *\*P<0.05*; *\*\*P<0.01*; *\*\*\*P<0.001*.

**Fig. S5 (related to Fig. 5).**
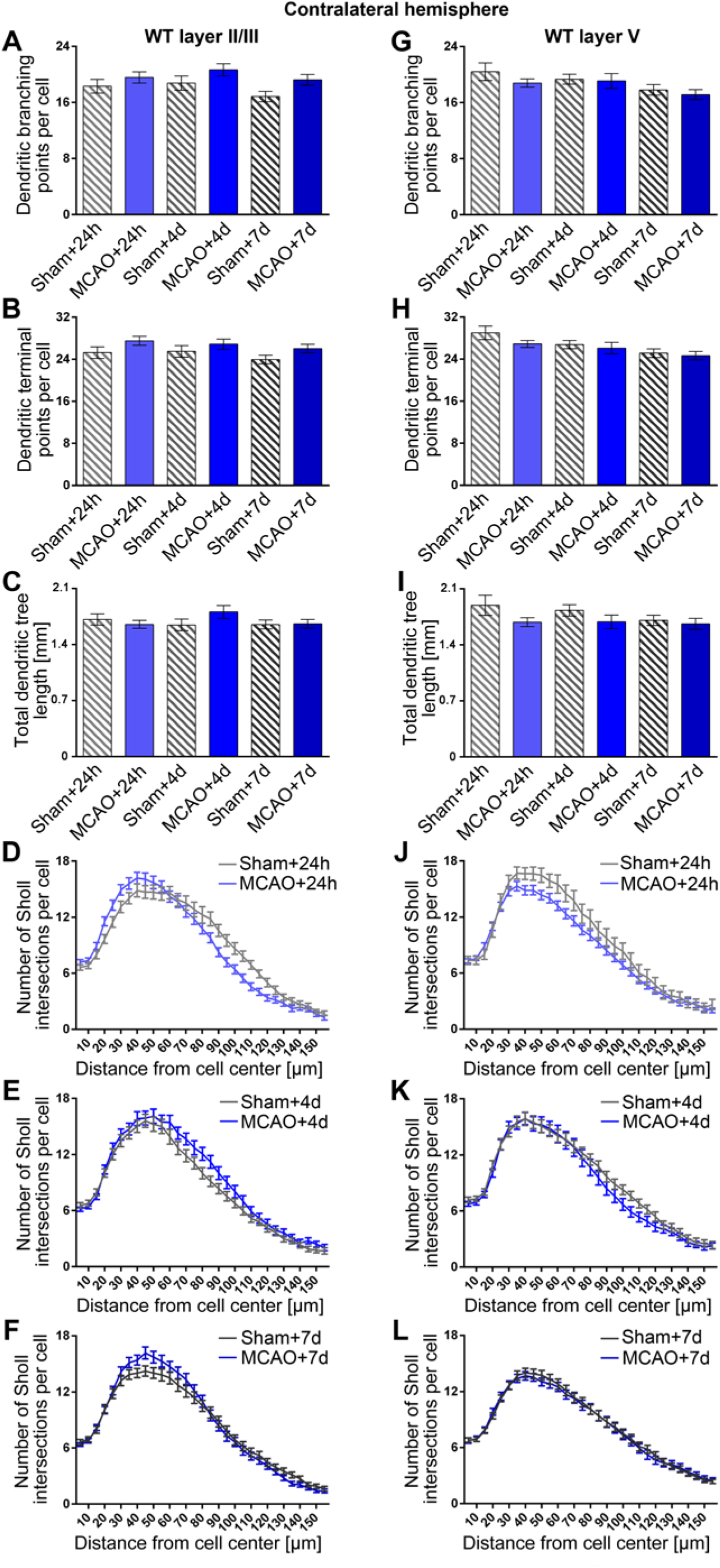
The dendritic arborization of neurons at the contralateral side is not affected by the MCAO-induced dendritic regrowth processes occurring simultaneously at the ipsilateral side. **A-L,** Quantitative determinations of dendritic arborization parameters of layer II/III neurons (**A-F**) and layer V neurons (**G-L**) in the contralateral M1. Note that, at the contralateral side, dendritic branching points (**A,G**), dendritic terminal points (**B,H**), total dendritic tree length (**C,I**) and Sholl intersections (**D-F,J-L**) at different reperfusion times all remained similar to their respective sham controls and at the same level for all three time points (24 h, 4 d, 7 d). Layer II/III: n_Sham+24h_=29; n_MCAO+24h_=53; n_Sham+4d_=38; n_MCAO+4d_=29; n_Sham+7d_=46; n_MCAO+7d_=49 neurons from 3 mice for the sham+24h, sham+4d and MCAO+4d groups and from 6 mice for the sham+7d, MCAO+24h and MCAO+7d groups (3-4 months of age). Layer V: n_Sham+24h_=17; n_MCAO+24h_=57; n_Sham+4d_=30; n_MCAO+4d_=30; n_Sham+7d_=57; n_MCAO+7d_=50 neurons from 3 mice of each group. For corresponding ipsilateral data see Fig. 5. Data, mean±SEM. Statistical significances were calculated using two-way ANOVA with Sidak’s post-test (all n.s.).

**Fig. S6 (related to Fig. 6).**
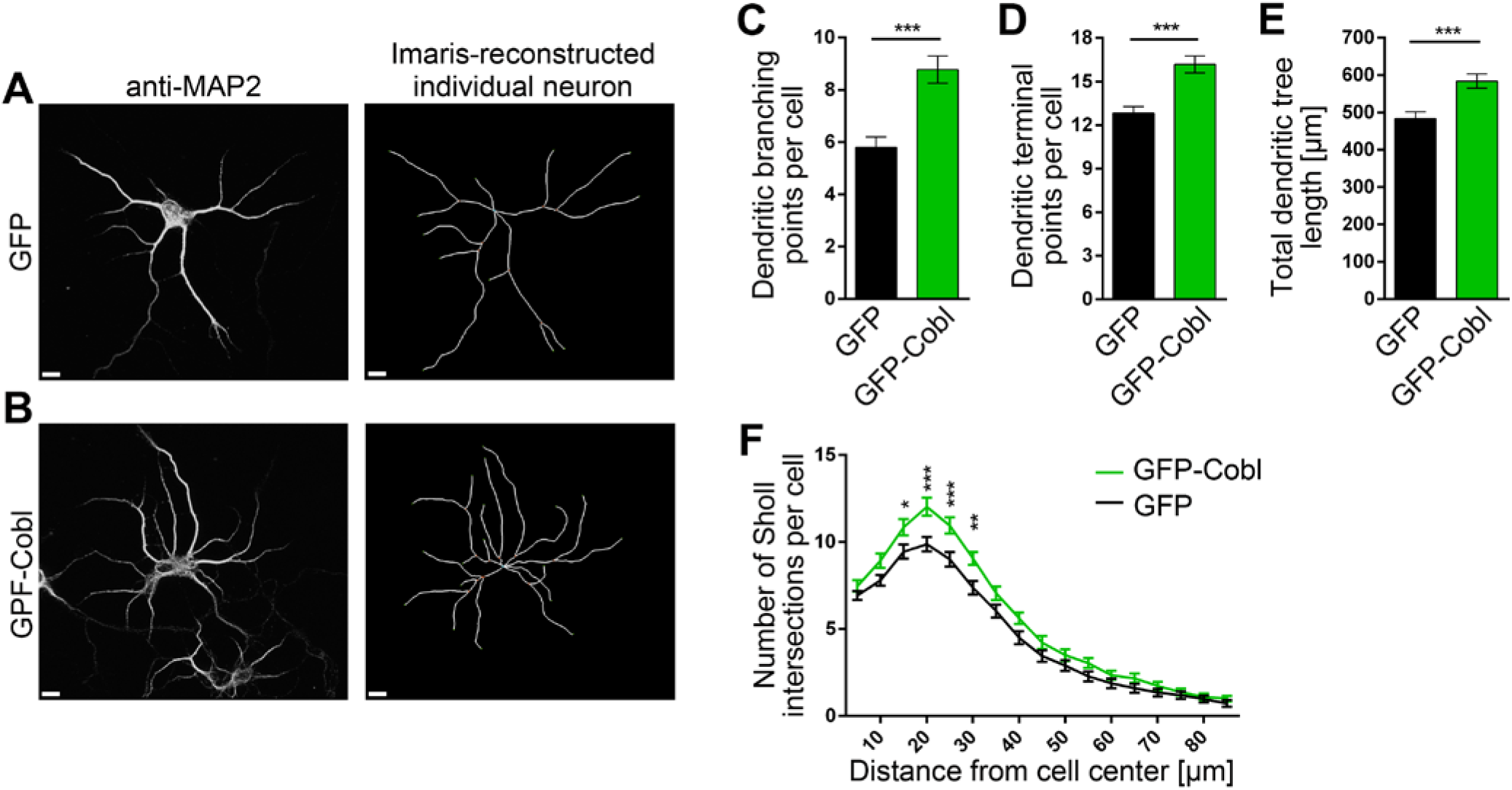
The actin nucleator Cobl promotes dendritic arborization during early development of cultured primary rat cortical neurons. **A,B,** Representative MIPs of anti-MAP2-immunostained (left panels) and Imaris-reconstructed primary rat cortical neurons (right panels) that were transfected at DIV4 with either GFP (**A**) and GFP-Cobl (**B**), respectively (marked by an asterisk), and fixed 40 h later. Scale bars, 10 μm. **C-F,** Quantitative determinations of dendritic arborization parameters unveiling clear Cobl gain-of-function phenotypes. Note that dendritic branching points (**C**), dendritic terminal points (**D**), total dendritic tree length (**E**) and Sholl intersections (**F**) all were increased upon Cobl overexpression, respectively. n_GFP_=40; n_GFP-Cobl_=40 individual transfected neurons from 2 independent preparations of cortical neurons. Data, mean±SEM. Statistical significances were calculated using Student’s t-test and two-way ANOVA with Sidak’s post-test for Sholl analysis. *\*P*< 0.05; *\*\*P*< 0.01; *\*\*\*P*< 0.001.

**Fig. S7 (related to Fig. 7).**
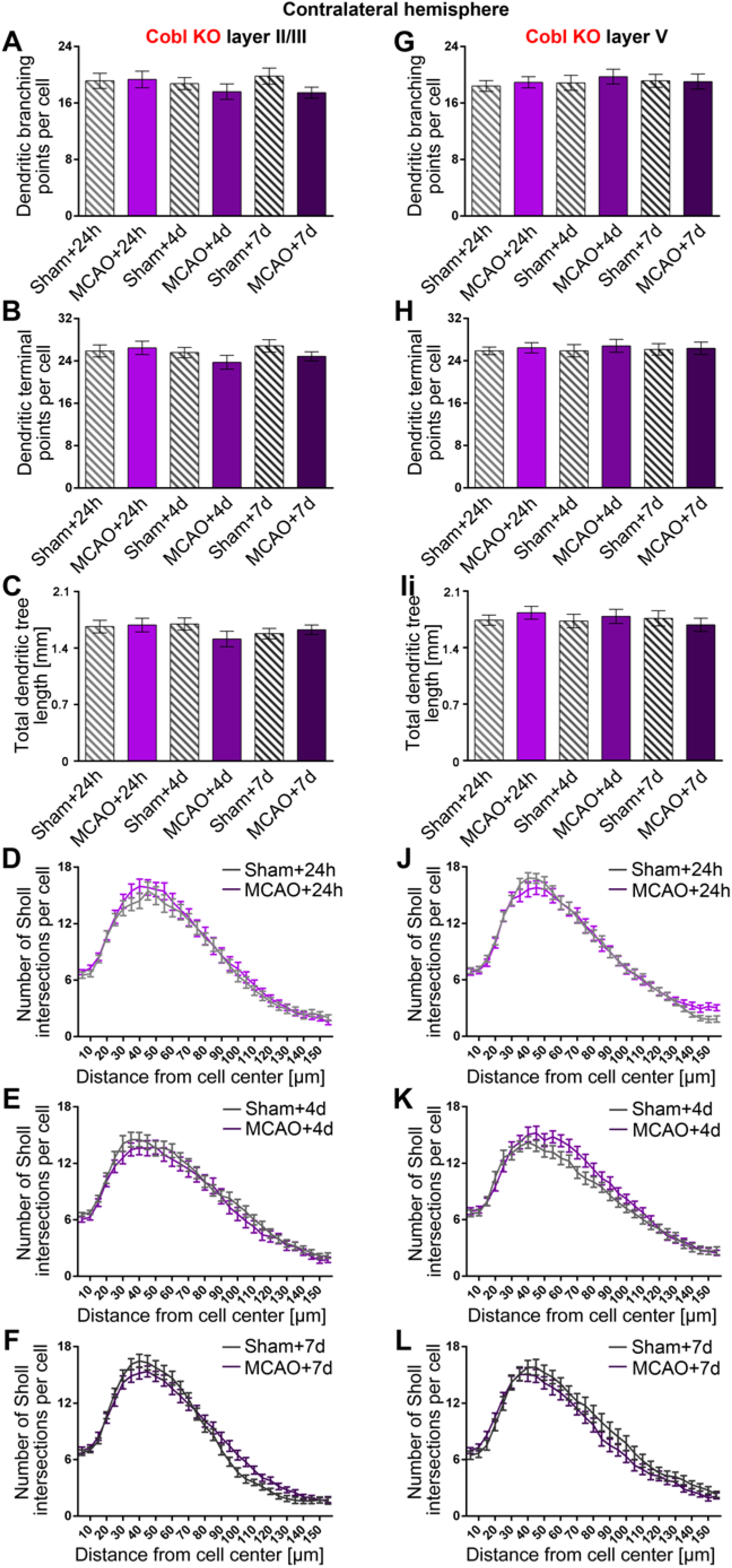
*Cobl* KO mice show no MCAO-induced dendritic alterations of the dendritic arbor at the contralateral side. **A-L,** Quantitative determinations of dendritic arborization parameters of layer II/III neurons (**A-F**) and layer V neurons (**G-L**) at the contralateral side in M1 of *Cobl* KO mice. Note that dendritic branching points (**A,G**), dendritic terminal points (**B,H**), total dendritic tree length (**C,I**) and Sholl intersections (**D-F,J-L**) at 24 h, 4 d and 7 d reperfusion times after 30 min MCAO at the juxtaposed side all remained unchanged and similar to their corresponding sham controls. Layer II/III: n_Sham+24h_=28; n_MCAO+24h_=32; n_Sham+4d_=33; n_MCAO+4d_=28; n_Sham+7d_=31; n_MCAO+7d_=36 neurons from 3 mice of each group. Layer V: n_Sham+24h_=34; n_MCAO+24h_=33; n_Sham+4d_=28; n_MCAO+4d_=34; n_Sham+7d_=26; n_MCAO+7d_=27 neurons from 3 mice of each group. For corresponding ipsilateral data see Fig. 7. Data represent mean±SEM. Statistical significances were calculated using two-way ANOVA with Sidak’s post-test (all n.s.).

**Fig. S8 (related to Fig. 7).**
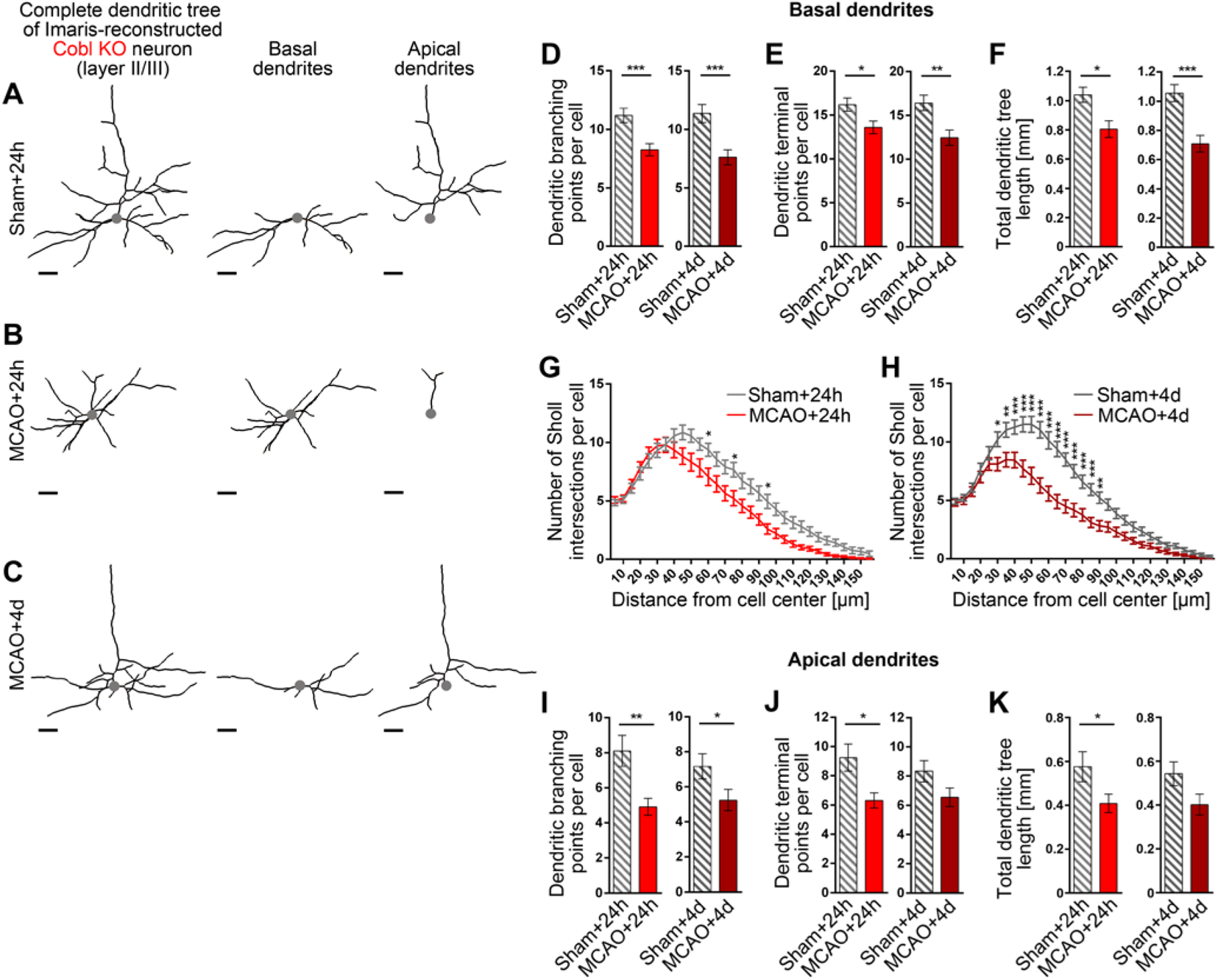
The defects in dendritic arbor repair observed in Cobl KO mice occur in both apical and basal dendrites of layer II/III pyramidal neurons of M1. **A-C,** Individual Imaris-reconstructed cells from Fig. 7A-C (right panels) showing ipsilateral layer II/III neurons from M1 of *Cobl* KO mice subjected to sham treatment (**A**) and 30 min MCAO with 24 h and 4 d reperfusion time (**B,C**), respectively, for differential analyses of basal (**D-H**) and apical dendritic parameters (**I-K**). The apical and basal dendritic parts are depicted in **A-C**. The somas are marked by a gray dot. Scale bars, 30 μm. **D-K,** Quantitative determinations of dendritic branching points, terminal points, total dendritic tree length and Sholl intersections of basal dendrites (**D-H**) and related analyses for apical dendrites (**I-K**). Note that MCAO-induced defects occurring in both apical and basal dendrites 24 after MCAO, in contrast to WT (see Fig. 5 and Fig. S5), do not show any recovery after 4 d reperfusion upon *Cobl* KO. n_Sham+24h_=21; n_MCAO+24h_=35; n_Sham+4d_=30; n_MCAO+4d_=29 neurons neurons from 3 mice for each MCAO and sham control group (ipsi). Quantitative data represent mean±SEM. Statistical significance calculations, Mann-Whitney (**D-F,I-K**) and two-way ANOVA with Sidak’s post-test for Sholl analysis (**E,F**), respectively. *\*P<0.05*; *\*\*P<0.01*; *\*\*\*P<0.001*.

## References

Ahuja, R., Pinyol, R., Reichenbach, N., Custer, L., Klingensmith, J., Kessels, M. M., and Qualmann, B. (2007). Cordon-bleu is an actin nucleation factor and controls neuronal morphology. Cell 131, 337–350.

Ankarcrona, M., Dypbukt, J. M., Bonfoco, E., Zhivotovsky, B., Orrenius, S., Lipton, S. A., and Nicotera, P. (1995). Glutamate-induced neuronal death: a succession of necrosis or apoptosis depending on mitochondrial function. Neuron 15, 961–973.

Beer, A. J., González Delgado, J., Steiniger, F., Qualmann, B., and Kessels, M. M. (2020). The actin nucleator Cobl organises the terminal web of enterocytes. Sci. Rep. 10, 11156.

Biernaskie, J., and Corbet, D. (2001). Enriched rehabilitative training promotes improved forelimb motor function and enhanced dendritic growth after focal ischemic injury. J. Neurosci. 21, 5272–5280.

Biernaskie, J., Chernenko, G., and Corbet, D. (2004). Efficacy of rehabilitative experience declines with time after focal ischemic brain injury. J. Neurosci. 24, 1245–1254.

Bogousslavsky, J., van Melle, G., and Regli, F. (1988). The Lausanne Stroke Registry: Analysis of 1,000 consecutive patients with first stroke. Stroke 19, 1083–1092.

Brown, C. E., Li, P., Boyd, J. D., Delaney, K. R., and Murphy, T. H. (2007). Extensive turnover of dendritic spines and vascular remodeling in cortical tissues recovering from stroke. J. Neurosci. 27, 4101–4109.

Brown, C. E., Wong, C., and Murphy, T. H. (2008). Rapid morphologic plasticity of peri-infarct dendritic spines after focal ischemic stroke. Stroke 39, 1286–1291.

Brown, C. E., Boyd, J. D., and Murphy, T. H. (2010). Longitudinal in vivo imaging reveals balanced and branch-specific remodeling of mature cortical pyramidal dendritic arbors after stroke. J. Cereb. Blood Flow Metab. 30, 783–791.

Calabrese, E. J. (2008). Neuroscience and hormesis: overview and general findings. Crit. Rev. Toxicol. 38, 249–252.

Corbett, D., Giles, T., Evans, S., McLean, J., and Biernaskie, J. (2006). Dynamic changes in CA1 dendritic spines associated with ischemic tolerance. Exp. Neurol. 202, 133–138.

Cramer, S. C, Shah, R., Juranek, J., Crafton, K. R., and Le, V. (2006). Activity in the peri-infarct rim in relation to recovery from stroke. Stroke 37, 111–115.

Das, G., Reuhl, K., and Zhou, R. (2013). The Golgi-Cox method. Methods Mol. Biol. 1018, 313–321.

Di Pino, G., Pellegrino, G., Assenza, G., Capone, F., Ferreri, F., Formica, D., Ranieri, F., Tombini, M., Ziemann, U., Rothwell, J. C., and Di Lazzaro, V. (2014). Modulation of brain plasticity in stroke: a novel model for neurorehabilitation. Nat. Rev. Neurol. 10, 597–608.

Fluri, F., Schuhmann, M. K., and Kleinschnitz, C. (2015). Animal models of ischemic stroke and their application in clinical research. Drug Des. Devel. Ther. 9, 3445–3454.

García-Chávez, D., González-Burgos, I., Letechipía-Vallejo, G., López-Loeza, E., Moralí, G., and Cervantes, M. (2008). Long-term evaluation of cytoarchitectonic characteristics of prefrontal cortex pyramidal neurons, following global cerebral ischemia and neuroprotective melatonin treatment, in rats. Neurosci. Lett. 448, 148–152.

Gascón, S., Sobrado, M., Roda, J. M., Rodríguez-Peña, A., and Díaz-Guerra, M. (2008). Excitotoxicity and focal cerebral ischemia induce truncation of the NR2A and NR2B subunits of the NMDA receptor and cleavage of the scaffolding protein PSD-95. Mol. Psychiatry 13, 99–114.

Glazner, G. W., Chan, S. L., Lu, C., and Mattson, M. P. (2000). Caspase-mediated degradation of AMPA receptor subunits: a mechanism for preventing excitotoxic necrosis and ensuring apoptosis. J. Neurosci. 20, 3641–3649.

Gonzalez, C.L.R., and Kolb, B. (2003). A comparison of different models of stroke on behaviour and brain morphology. Eur. J. Neurosci. 18, 1950–1962.

Graham, F. L., Smiley, J., Russell, W. C., Nairn, R. (1977). Characteristics of a human cell line transformed by DNA from human adenovirus type 5. J. Gen. Virol. 36, 59–74.

Haag, N., Schüler, S., Nietzsche, S., Hübner, C. A., Strenzke, N., Qualmann, B., and Kessels, M. M. (2018). The actin nucleator Cobl is critical for centriolar positioning, postnatal planar cell polarity refinement, and function of the cochlea. Cell Rep. 24, 2418–2431.

Haag, N., Schwintzer, L., Ahuja, R., Koch, N., Grimm, J., Heuer, H., Qualmann, B., and Kessels, M. M. (2012). The actin nucleator Cobl is crucial for Purkinje cell development and works in close conjunction with the F-actin binding protein Abp1. J. Neurosci. 32, 17842–17856.

Haeckel, A., Ahuja, R., Gundelfinger, E. D., Qualmann, B., and Kessels, M. M. (2008). The actin-binding protein Abp1 controls dendritic spine morphology and is important for spine head and synapse formation. J. Neurosci. 28, 10031–10044.

Heiss, W. D. (2000). Ischemic penumbra: Evidence from functional imaging in man. J. Cereb. Blood Flow Metab. 20, 1276–1293.

Hossmann, K. A. (2006). Pathophysiology and therapy of experimental stroke. Cell. Mol. Neurobiol. 26, 1057–1083.

Hou, W., Izadi, M., Nemitz, S., Haag, N., Kessels, M. M., and Qualmann, B. (2015). The actin nucleator Cobl is controlled by calcium and calmodulin. PLoS Biol. 13, e1002233.

Hou, W., Nemitz, S., Schopper, S., Nielsen, M. L., Kessels, M. M., and Qualmann, B. (2018). Arginine methylation by PRMT2 controls the functions of the actin nucleator Cobl. Dev. Cell 45, 262–275.

Hu, J., Li C., Hua, Y., Liu, P., Gao, B., Wang, Y., and Bai, Y. (2020). Constraint-induced movement therapy improves functional recovery after ischemic stroke and its impacts on synaptic plasticity in sensorimotor cortex and hippocampus. Brain Res. Bull. 160, 8–23.

Izadi, M., Schlobinski, D., Lahr, M., Schwintzer, L., Qualmann, B., and Kessels, M. M. (2018). Cobl-like promotes actin filament formation and dendritic branching using only a single WH2 domain. J. Cell Biol. 217, 211–230.

Izadi, M., Seemann, E., Schlobinski D., Schwintzer, L., Qualmann, B., and Kessels, M. M. (2021). Functional interdependence of the actin nucleator Cobl and Cobl-like in dendritic arbor development. eLIFE, 10, e67718.

Jaenisch, N., Liebmann, L., Guenther, M., Hübner, C. A., Frahm, C., and Witte, O. W. (2016). Reduced tonic inhibition after stroke promotes motor performance and epileptic seizures. Sci Rep. 6, 26173.

Jones, T.A. (2017). Motor compensation and its effects on neural reorganization after stroke. Nat. Rev. Neurosci. 18, 267–280.

Kessels, M. M., Schwintzer, L., Schlobinski, D., and Qualmann, B. (2011). Controlling actin cytoskeletal organization and dynamics during neuronal morphogenesis. Eur. J. Cell Biol. 90, 926–933.

Kim, Y., Sung, J. Y., Ceglia, I., Lee, K. -W., Ahn, J. -H., Halford, J. M., Kim, A. M., Kwak, S. P., Park, J. B., Ryu, S. H., Schenck, A., Bardoni, B., Scott, J. D., Nairn, A. C., and Greengard, P., (2006). Phosphorylation of WAVE1 regulates actin polymerization and dendritic spine morphology. Nature 442, 814–817.

Koch N., Koch, D., Krueger, S., Tröger, J., Sabanov, V., Ahmed, T., McMillan, L. E., Wolf, D., Montag, D., Kessels, M. M., Balschun, D., and Qualmann, B. (2020). Syndapin I loss-of-function in mice leads to schizophrenia-like symptoms. Cereb. Cortex. 30, 4306–4324.

Koleske, A. J. (2013). Molecular mechanisms of dendrite stability. Nat. Rev. Neurosci. 14, 536–550.

Murphy, T. H. and Corbett, D. (2009). Plasticity during stroke recovery: from synapse to behavior. Nat. Rev. Neurosci. 10, 861–872.

Lynch, G., and Baudry, M. (1984). The biochemistry of memory: A new and specific hypothesis. Science 224, 1057–1063.

Mattson, M.P., and Cheng, A. (2006). Neurohormetic phytochemicals: Low-dose toxins that induce adaptive neuronal stress responses. Trends Neurosci. 29, 632–639.

Mauceri, D., Buchthal, B., Hemstedt, T. J., Weiss, U., Klein, C. D., and Bading, H. (2020). Nasally delivered VEGFD mimetics mitigate stroke-induced dendrite loss and brain damage. Proc. Natl. Acad. Sci. USA, 117, 8616–8623.

Mostany, R., and Portera-Cailliau, C. (2011). Absence of large-scale dendritic plasticity of layer 5 pyramidal neurons in peri-infarct cortex. J. Neurosci. 31, 1734–1738.

Musuka, T. D., Wilton, S. B., Traboulsi, M., and Hill, M. D. (2015). Diagnosis and management of acute ischemic stroke: speed is critical. CMAJ 187, 887–893.

Nesin, S. M., Sabitha, K. R., Gupta, A., and Laxmi, T. R. (2019). Constraint induced movement therapy as a rehabilitative strategy for ischemic stroke-linking neural plasticity with restoration of skilled movements. J. Stroke Cerebrovasc. Dis. 28, 1640–1653.

Nudo, R. J., Wise, B. M., SiFuentes, F., and Milliken, G. W. (1996). Neural substrates for the effects of rehabilitative training on motor recovery after ischemic infarct. Science 272, 1791–1794.

Papadopoulos, C. M., Tsai, S., Cheatwood, J. L., Bollnow, M. R., Kolb, B. E., Schwab, M. E., and Kartje, G. L. (2006). Dendritic plasticity in the adult rat following middle cerebral artery occlusion and Nogo-a neutralization. Cereb. Cortex. 16, 529–536.

Pfaffl, M. W. (2001). A new mathematical model for relative quantification in real-time RT-PCR. Nucleic Acids Res. 29, e45.

Pike, B. R., Flint, J., Dave, J. R., Lu, X. M., Wang, K. K. K., Tortella, F. C., and Hayes, R. L. (2004). Accumulation of calpain and caspase-3 proteolytic fragments of brain-derived αII-Spectrin in cerebral spinal fluid after middle cerebral artery occlusion in rats. J. Cereb. Blood Flow Metab. 24, 98–106.

Pizarro-Cerdá, J., Chorev, D. S., Geiger, B., and Cossart, P. (2016). The diverse family of Arp2/3 complexes. Trends Cell Biol. 27, 93–100.

Popp, A., Jaenisch, N., Witte, O.W., and Frahm, C. (2009). Identification of ischemic regions in a rat model of stroke. PLoS One. 4, e4764.

Qualmann, B. and Kessels, M. M. (2009). New players in actin polymerization ‒ WH2-domain-containing actin nucleators. Trends Cell Biol. 19, 276–285.

Ravanelli, A. M., and Klingensmith, J. (2011). The actin nucleator Cordon-bleu is required for development of motile cilia in zebrafish. Dev Biol. 350, 101–111.

Schneider, K., Seemann, E., Liebmann, L., Ahuja, R., Koch, D., Westermann, M., Hübner, C. A., Kessels, M. M., and Qualmann, B. (2014). ProSAP1 and membrane nanodomain-associated syndapin I promote postsynapse formation and function. J. Cell Biol. 205, 197–215.

Schüler, S., Hauptmann, J., Perner, B., Kessels, M. M., Englert, C., and Qualmann, B. (2013). Ciliated sensory hair cell formation and function require the F-BAR protein syndapin I and the WH2 domain-based actin nucleator Cobl. J. Cell Sci. 126, 196–208.

Schwintzer, L., Koch, N., Ahuja, R., Grimm, J., Kessels, M. M., and Qualmann, B. (2011). The functions of the actin nucleator Cobl in cellular morphogenesis critically depend on syndapin I. EMBO J. 30, 3147–3159.

Sieber, M. W., Guenther, M., Kohl, M., Witte, O. W., Claus, R. A., and Frahm, C. (2010) Inter-age variability of bona fide unvaried transcripts normalization of quantitative PCR data in ischemic stroke. Neurobiol. Aging 31, 654–664.

Siman, R., Baudry, M., and Lynch, G. (1984). Brain fodrin: substrate for calpain I, an endogenous calcium-activated protease. Proc. Natl. Acad. Sci. U. S. A. 81, 3572–3576.

Simpkins, K. L., Guttmann, R. P., Dong, Y., Chen, Z., Sokol, S., Neumar, R. W., and Lynch, D. R. (2003). Selective activation induced cleavage of the NR2B subunit by calpain. J. Neurosci. 23, 11322–11331.

Soderling, S. H., Guire, E. S., Kaech, S., White, J., Zhang, F., Schutz, K., Langeberg, L. K., Banker, G., Raber, J., and Scott, J. D. (2007). A WAVE-1 and WRP signaling complex regulates spine density, synaptic plasticity, and memory. J. Neurosci. 27, 355–365.

Spence, E. F., Kanak, D. J., Carlson, B. R., and Soderling, S. H. (2016). The Arp2/3 complex is essential for distinct stages of spine synapse maturation, including synapse unsilencing. J. Neurosci. 36, 9696–709.

Stahr, A., Frahm, C., Kretz, A., Bondeva, T., Witte, O. W., and Wolf, G. (2012). Morg1(+/-) heterozygous mice are protected from experimentally induced focal cerebral ischemia. Brain Res. 1482, 22–31.

Wegner, A. M., Nebhan, C. A., Hu, L., Majumdar, D., Meier, K. M., Weaver, A. M., and Webb, D. J. (2008). N-WASP and the Arp2/3 complex are critical regulators of actin in the development of dendritic spines and synapses. J. Biol. Chem. 283, 15912–15920.

Winship, I. R. and Murphy, T. H. (2008). In vivo calcium imaging reveals functional rewiring of single somatosensory neurons after stroke. J. Neurosci. 28, 6592–6606.

Wolf, D., Hofbrucker-MacKenzie, S. A., Izadi, M., Seemann, E., Steiniger, F., Schwintzer, L., Koch, D., Kessels, M. M., and Qualmann, B. (2019). Ankyrin repeat-containing N-Ank proteins shape cellular membranes. Nat. Cell Biol. 21, 1191–1205.

Wu, F., Catano, M., Echeverry, R., Torre, E., Haile, W. B., An, J., Chen, C., Cheng, L., Nicholson, A., Tong, F. C., Park, J., and Yepes, M. (2014). Urokinase-type plasminogen activator promotes dendritic spine recovery and improves neurological outcome following ischemic stroke. J. Neurosci. 34, 14219–14232.

Xu, J., Kurup, P., Zhang, Y., Goebel-Goody, S. M., Wu, P. H, Hawasli, A. H, Baum, M. L, Bibb, J. A., and Lombroso, P. L. (2009). Extrasynaptic NMDA receptors couple preferentially to excitotoxicity via calpain-mediated cleavage of STEP. J. Neurosci. 29, 9330–9343.

